# Rotten to the core – a neurofunctional signature of subjective core disgust generalizes to oral distaste and socio-moral contexts

**DOI:** 10.1101/2023.05.18.541259

**Authors:** Xianyang Gan, Feng Zhou, Ting Xu, Xiaobo Liu, Ran Zhang, Zihao Zheng, Xi Yang, Xinqi Zhou, Fangwen Yu, Jialin Li, Ruifang Cui, Lan Wang, Jiajin Yuan, Dezhong Yao, Benjamin Becker

## Abstract

While disgust originates in the hard-wired mammalian distaste response, the conscious experience of disgust in humans strongly depends on subjective appraisal and may even extend to sociomoral contexts. In a series of studies, we combined functional magnetic resonance imaging (fMRI) with machine-learning based predictive modeling to establish a comprehensive neurobiological model of subjective disgust. The developed neurofunctional signature accurately predicted momentary self-reported subjective disgust across discovery (*n*=78) and pre-registered validation (*n*=30) cohorts and generalized across core disgust (*n*=34 and *n*=26), gustatory distaste (*n=30*), and sociomoral (unfair offers; *n*=43) contexts. Disgust experience was encoded in distributed cortical and subcortical systems, and exhibited distinct and shared neural representations with subjective fear or negative affect in interoceptive-emotional awareness and conscious appraisal systems while the signatures most accurately predicted the respective target experience. We provide an accurate fMRI-signature for disgust with a high potential to resolve ongoing evolutionary debates.

## Introduction

Across cultures, disgust is commonly triggered by rotting food, maggots, and scavenging animals but also by highly culture- and individual-specific triggers such as specific foods or behaviours that are appraised as morally reprehensible. Disgust has been conceptualized as a multifaceted defensive-avoidance response that encompasses hard-wired physiological and motor responses such as retching or a distinct facial expression and a strongly aversive feeling of revulsion that together facilitate the avoidance of potentially poisonous, contagious, or morally offensive situations and individuals. From an evolutionary perspective, disgust originates in the mammalian bitter taste (distaste) rejection system aiming to protect against the ingestion of potentially toxic food^1–4^. In response to bitter food, the distaste mechanism directly triggers defensive reactions such as withdrawal and retching and distinct facial expressions across (some) mammalian species via brainstem circuits^5^ or the posterior insular cortex^6^, respectively (for the disgust output system^1^ see also Fig. 1a). In humans, the distaste response was preadapted during evolution into core disgust which is essentially characterized by the highly aversive feeling of revulsion in response to contaminated, infectious, or poisonous stimuli, in turn facilitating pathogen avoidance^7–10^ (thus also termed as pathogen disgust according to Tybur et al.^11,12^). Core disgust is distinct from the distaste reaction given that the experience of revulsion can be evoked in the absence of actual sensory stimuli like bitter taste and is strongly shaped by conscious appraisals (e.g., food being rejected based on where it might have been)^10^. Subsequently, core disgust may have been preadapted into the social context with revulsion being elicited by sociomoral transgressions^1,2,4,13^ which may facilitate avoidance of potentially harmful interactions (sociomoral disgust)^14^. The preadaptation processes of disgust described above are further reflected in and supported by adaptationist perspectives^11,12^ that posit that disgust has evolved in response to pathogen avoidance and was subsequently co-opted to serve adaptive purposes in the realm of sociomoral disgust.

**Fig. 1.**
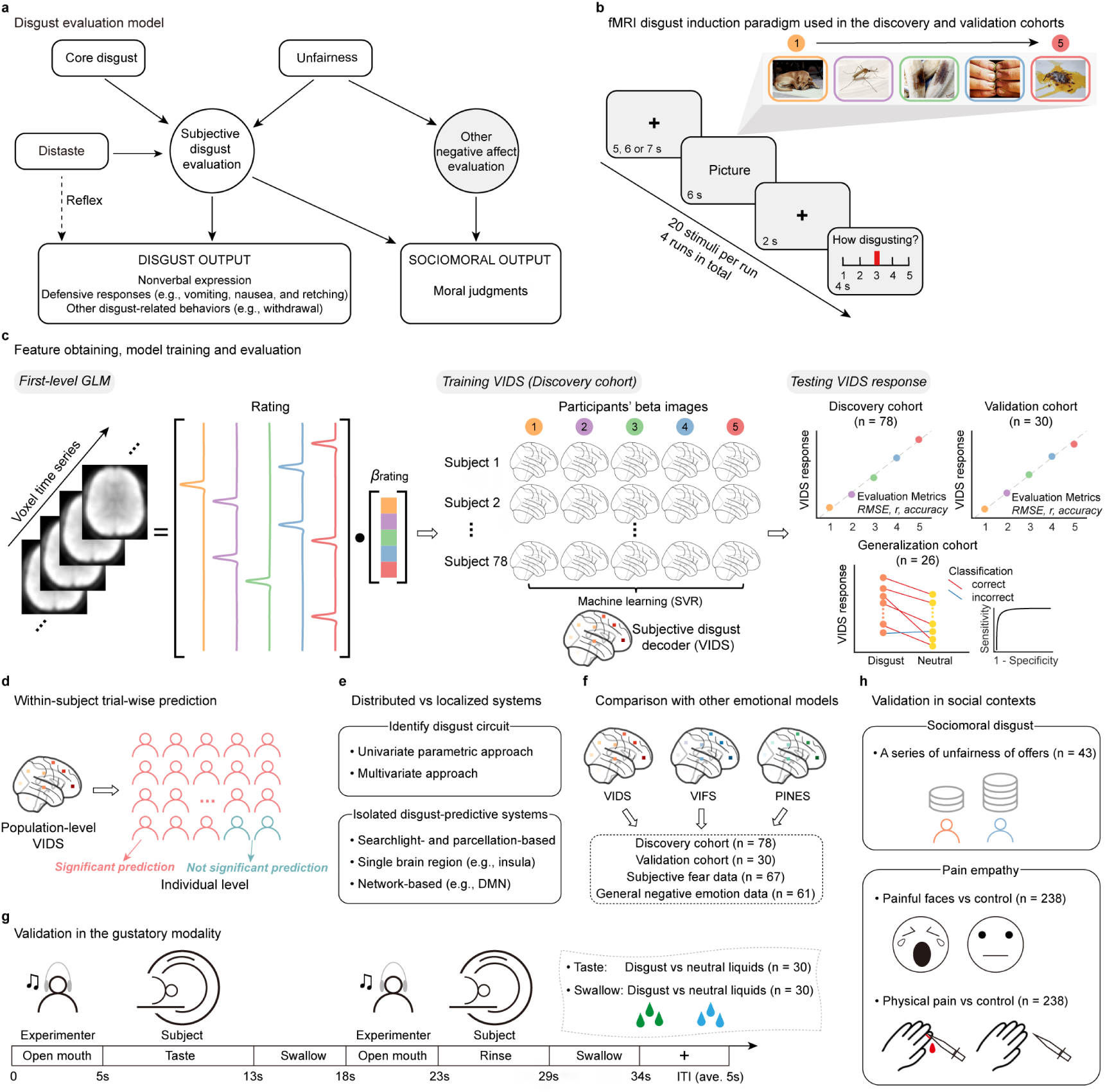
Disgust evaluation model, task design, and analytic workflow. **a**, Disgust evaluation model proposed based on previous studies^1,14^. **b**, Discovery and validation cohorts underwent the identical disgust induction fMRI paradigm. Example stimuli from the DIRTI disgust images database^59^ (note that all pictures from the DIRTI are subject to creative commons license CC BY-NC 4.0). **c**, Feature determination, model training, and evaluation. Voxel-level brain maps (beta images) were used as features in the prediction analysis. A whole-brain multivariate pattern predictive of the level of subjective disgust (VIDS) was trained on the discovery sample (*n*=78) using SVR and further evaluated in discovery (cross-validated), validation (*n*=30), and generalization (*n*=26) cohorts. **d**, Within-subject trial-wise prediction by applying the population-level VIDS to individual level trial-wise disgust experience in both discovery and validation cohorts. **e**, Systematic tests of distributed versus localized disgust systems hypotheses. Univariate and multivariate approaches were employed to determine the contribution of specific brain systems to predict subjective disgust. Multiple prediction analyses were next conducted to test the performance of isolated brain regions or systems in predicting subjective disgust experience. **f**, Testing the specificity of the VIDS in terms of distinguishable representations compared to subjective fear or general negative affect. **g**, Validation of disgust in the gustatory modality. The experimental flowchart depicts the gustatory procedures. The experimenter receiving auditory instructions was responsible for the delivery of gustatory stimuli, the subject (lying supine) followed the experimental procedures by looking at the visual instructions shown on the screen. For each trial, 1ml of liquid (either disgust or neutral taste) was delivered in the mouth, which had to be held in the mouth and tasted for 8s, followed by a swallow instruction (5s). To avoid transfer of the remaining taste to the next trial, 2ml of water was then delivered in the mouth, which was used for rinse (6s), again followed by a 5s swallow instruction. Inter-trial intervals (ITI) ranged from 4-6s (average 5s). Next, validations in the gustatory modality were performed for both the taste and swallow of disgust versus neutral liquids (the dotted region at the upper right panel). **h**, Validation of disgust in social contexts. The VIDS was applied to two social contexts, i.e., sociomoral disgust (unfairness), and pain empathy induced by physical (limb) and affective (face) pain-displaying stimuli. DMN=default mode network; GLM=general linear model; RMSE=root mean squared error; SVR=support vector regression; VIDS=visually-induced disgust signature developed in this study; VIFS=visually-induced fear signature from Zhou et al.^18^; PINES=picture-induced negative emotion signature from Chang et al.^39^. See Methods and Supplementary Table 1 for further details.

While several neurobiological models on the exact brain mechanisms of the distaste reaction have been established in animal models^5,15,16^, recent conceptual and computational advances indicate that the neural mechanisms that underlie conscious emotional experiences are distinguishable from those that mediate the hard-wired defensive and social-communicative facets of the response^8,17–21^. While these studies primarily focused on fear, the neural mechanisms underlying the conscious experience of disgust, whether these mechanisms differ from the experience of evolutionary similar defensive functions (e.g., fear), and whether the same mechanisms mediate the aversive experience in the gustatory distaste response or in sociomoral transgressions remain fiercely debated.

A plethora of human studies employed functional Magnetic Resonance Imaging (fMRI) to determine the neural basis of disgust, yet findings with respect to the neural systems that mediate conscious disgust experience remained highly inconsistent. For example, the insula has been traditionally considered as a key neural substrate of disgust^7,8,22,23^, however several studies did not observe insula engagement in response to disgust stimuli^24–28^ and recent evidence suggests a broader role of the insula in interoceptive^29,30^, central autonomic^31^, and individual-level emotional information processing^32,33^. Subcortical regions like the amygdala and basal ganglia have also been considered as key disgust regions^8,22,34^, probably due to their roles in early threat detection and defensive motor responses^8,17,18^. However, many neuroimaging studies failed to observe an involvement of amygdala^26–28,35–37^ while a recent meta-analysis did not confirm robust engagement of the basal ganglia^8^ during disgust processing. These inconsistencies may arise via a combination of (1) experimental designs that do not account for individual differences in disgust appraisal and experience^38^ and the employed categorical analytic approach of comparing highly disgusting versus neutral stimuli which inherently also captures neural activity associated with the defensive-avoidance response, salience or arousal^39^, and, (2) methodological limitations of the conventional mass-univariate fMRI approach with respect to establishing accurate, comprehensive and process-specific neural models for mental processes^40,41^.

Recent methodological progress combined fMRI with machine-learning based multivariate pattern analysis (MVPA) to develop more comprehensive and accurate neurofunctional models of internal mental processes, including sensitive and generalizable neuroaffective signatures for subjective emotional experiences (e.g., negative affect or fear)^18,39,42,43^ as compared to conventional brain imaging models. Identifying multivariate brain signatures predictive of the momentary emotional experience may allow to overcome critical limitations of the mass-univariate fMRI approach while accounting for contextual and individual variations on the appraisal and experience level^44^. Briefly, MVPA examines neural responses across multiple brain regions and voxels^40,45–47^ thus allowing to determine neural representations of mental processes with higher resolution and considerably larger effect sizes^40,48^ than the conventional mass-univariate approach. The resultant fMRI-based neurofunctional signatures enabled for the first time to separate the subjective emotional experience from hard-wired defensive and physiological responses^18,20^ and to explore common and separable neural representations of aversive experiences across domains^43,49^. In line with recent conceptual frameworks, these findings indicate that the concerted interplay between subcortical and cortical systems mediates the appraisal and construction of conscious emotional experiences^50,51^. Despite long-standing discussions on the neurobiological and cultural evolution of disgust and increasing interest in the role of disgust in a number of mental and neurological (e.g., obsessive-compulsive disorder, eating disorders, and Huntington’s disease)^7,9,34^, and initial studies demonstrated the promise of MVPA for regional or categorical disgust decoding^52,53^, an accurate and generalizable whole-brain signature predictive of momentary self-reported subjective disgust experience is currently lacking.

Here, we capitalized on recent methodological advances in MVPA-based neural decoding techniques to determine (1) whether it is possible to develop a neural signature that predicts momentary subjective disgust experience elicited by core disgust stimuli on the population-level with high sensitivity and robustness across a discovery (*n*=78 healthy participants) and a pre-registered validation (*n*=30) cohort, that generalizes to data from an independent research team using a different disgust induction paradigm and MRI system (*n*=26)^54^ (Fig. 1b,c) as well as a modified disgust induction paradigm that uncouples neural activity related to the emotional experience and motor responses (*n*=34); (2) whether this neural signature can sensitively track variations in momentary disgust experience on the individual level in terms of predicting trial-wise disgust experience in discovery and validation cohorts (Fig. 1d); (3) the neural representations underlying disgust experience utilizing forward (association), backward (prediction) encoding models^55^, and isolated systems predictions (notably, the MVPA approach provides the opportunity to test the performance of local systems versus distributed whole-brain activity pattern^56^) (Fig. 1e). The developed visually-induced disgust signature (VIDS) was next utilized to determine (4) the functional specificity of the subjective disgust signature by conducting predictive and topographical comparisons with established decoders for subjective fear^18^ and negative affect^39^, respectively (Fig. 1f); (5) whether the visual disgust decoder generalizes to the gustatory modality by applying the VIDS to data acquired while participants tasted (as well as swallowed) either disgust or neutral liquids (*n=30*) (Fig. 1g); (6) whether core disgust also extends to social experiences by applying the VIDS to data from two experiments conducted in social contexts, including neural reactivity in response to unfair offers in an Ultimatum Game (sociomoral disgust, *n*=43) and in response to the suffering of other individuals in a pain empathy paradigm^49^ (*n*=238) (Fig. 1h). Finally, control analyses were employed in two additional independent datasets that examined neural reactivity in response to positive and negative visual emotional stimuli^57,58^ (*n*=30, *n*=150, respectively) to explore the extent to which the VIDS may capture unspecific emotional arousal. Together, this systematic examination enables us to establish a comprehensive brain-based model for subjective disgust experience and to determine the extent of neurofunctional similarity with the experience of other negative emotions (fear and negative affect), as well as with the experience of distaste in the gustatory modality and unfairness in social situations (commonly referred to as sociomoral disgust) hypothesized in the disgust evaluation model (Fig. 1a).

## Results

### Visual disgust stimuli robustly evoked different levels of subjective disgust experience

During fMRI, disgust was induced by means of validated disgust-specific affective pictures^59^ (DIRTI disgust images database, details see Methods). Participants were instructed to react naturally to the stimuli and report their momentary level of disgust during each trial on a 5-point Likert scale ranging from 1 (neutral/slightest disgust) to 5 (very strong disgust). 80 stimuli were presented and robustly induced the entire intensity range of subjective disgust balanced across the most common disgust-related contexts (animal, human, and scene; Extended Data Fig. 1). In the discovery cohort (*n*=78) utilized to develop the disgust signature, the stimuli evoked sufficient levels of subjective disgust experience across contexts such that over 12% of stimuli in each context induced very strong disgust (rating 5), with 72 out of 78 participants reporting all 5 levels of subjective disgust (6 reported levels 1-4).

### A brain signature sensitive to predict subjective disgust experience

Consistent with previous studies developing neuro-affective decoders^18,20,39^, we employed a linear support vector regression (SVR) model to identify a whole-brain signature of fMRI activation predictive of subjective disgust experience (Fig. 2a and Extended Data Fig. 2). Notably, previous studies (e.g., refs.^39,60^) have shown that linear SVR exhibits nearly identical prediction performance to linear models with regulations and feature reduction. The decoder was developed in the discovery cohort, next performance of the VIDS was evaluated using cross-validation in the discovery cohort (10×10-fold cross-validated, Methods) and by determining reactivity of the VIDS to disgust experience in an independent and pre-registered validation cohort (*n*=30) that underwent the identical paradigm. The developed VIDS accurately predicted the level of subjective disgust in the discovery cohort (averaged within-subject correlation between predicted and true disgust ratings, 5 or 4 pairs of scalar values per subject: *r*=0.88±0.02 (mean±standard error, SE); averaged RMSE=1.43±0.08; overall, i.e., between- and within-subject, prediction-outcome correlation for 384 pairs, *r*=0.56, averaged across ten repetitions, bootstrapped 95% confidence interval (CI)=[0.49, 0.62], Fig. 2b). To independently determine the robustness of the VIDS, we pre-registered an independent validation cohort (https://osf.io/y4s78). Applying the predictive model - with no further model fitting – to the validation cohort revealed comparable high prediction-outcome correlations (within-subject *r*=0.89±0.03, average RMSE=1.21±0.14; overall prediction-outcome *r*=0.62, 95% CI=[0.53, 0.70]; Fig. 2c), indicating a sensitive and robust neurofunctional subjective disgust signature. To further determine the sensitivity of the model for subjective disgust, we compared pairs of activation maps within each subject by means of a two-alternative forced-choice test choosing the activation map with higher VIDS response as the one with higher disgust experience. The VIDS response accurately classified high (average of rating 4 and 5) versus moderate (rating 3) and moderate versus low (average of rating 1 and 2) disgust in both cohorts with 86%-96% accuracy (Cohen’s *d*=1.22–1.96), and high versus low with 99%-100% accuracy in both cohorts (Cohen’s *d*=2.40–2.92). Moreover, the VIDS response could distinguish each successive pair of disgust rating levels (e.g., rating 1 versus rating 2) with at least 80% accuracy (significantly higher than 50% chance-level; *P*<0.001; except rating 5 versus 4 in the validation cohort, with accuracy being 76% and *P*<0.01) (Fig. 2b,c).

**Fig. 2.**
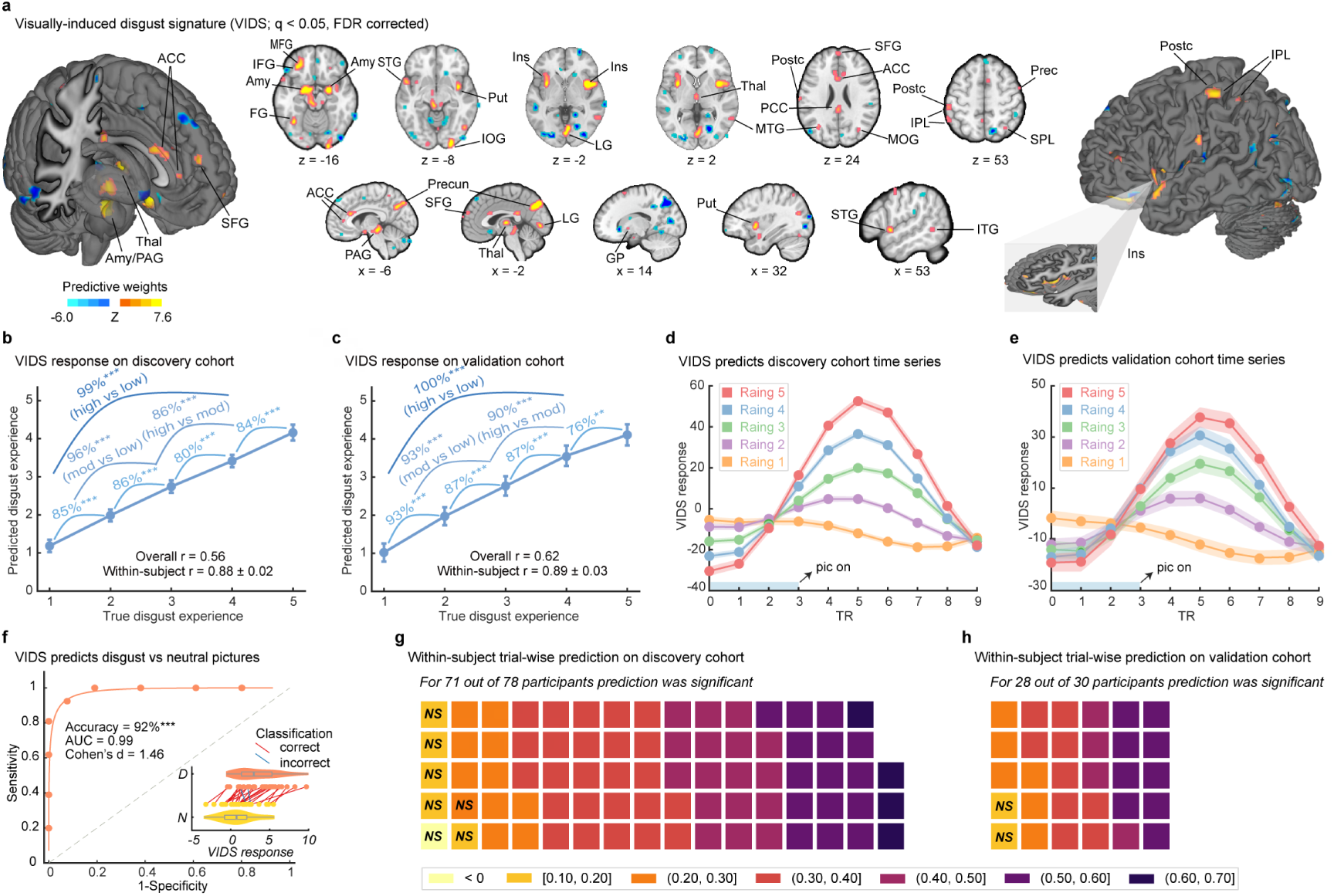
Visually-induced disgust signature (VIDS). **a**, The VIDS pattern thresholded at *q*<0.05 (FDR corrected, 10,000-sample bootstrap). Extended Data Fig. 2 shows the topography of the unthresholded pattern in anatomical regions of interest previously associated with disgust and emotional experience. **b,c**, Predicted disgust experience (subjective ratings; mean±SE) compared to actual disgust ratings in the cross-validated discovery cohort (*n*=78) and the independent validation cohort (*n*=30), respectively. Accuracy provided for forced-choice comparisons. *P* values based on two-sided independent binomial tests. *r* indicates Pearson correlation coefficient between predicted and true ratings. **d,e**, Averaged peristimulus plot (mean±SE) of the VIDS response in the cross-validated discovery cohort and the independent validation cohort, respectively, at every repetition time (TR; 2s) for each disgust intensity rating separately. **f**, Generalization of the VIDS in an independent dataset (using two-alternative forced-choice to classify disgust vs neutral stimuli). The figure shows the receiver-operating characteristics (ROC), and the inlet shows the VIDS response for different conditions (D=disgust; N=neutral). Each line connecting dots represents paired data from the same participant (red=correct classification; blue=incorrect classification). *P* value was based on a binomial test, two-sided. **g,h**, Range of the within-subject trial-wise prediction-outcome correlation coefficients for each participant in discovery and validation cohorts, respectively; note that the grids represent the prediction performance on the individual level, with each cell representing prediction performance (in terms of prediction-outcome correlation) for an individual participant. ** *P*<0.01, *** *P*<0.001, NS not significant. Error bars and shaded regions indicate SEs. ACC=anterior cingulate cortex; Amy=amygdala; AUC=area under the curve; FDR=false discovery rate; FG=fusiform gyrus; GP=globus pallidus; IFG=inferior frontal gyrus; Ins=insula; IOG=inferior occipital gyrus; IPL=inferior parietal lobule; ITG=inferior temporal gyrus; LG=lingual gyrus; MFG=middle frontal gyrus; MOG=middle occipital gyrus; MTG=middle temporal gyrus; PAG=periaqueductal gray; PCC=posterior cingulate cortex; Postc=postcentral gyrus; Prec=precentral gyrus; Precun=precuneus; Put=putamen; SFG=superior frontal gyrus; SPL=superior parietal lobule; STG=superior temporal gyrus; Thal=thalamus.

Given that previous studies emphasized the contribution of the visual cortex in decoding emotional content^8,61^, we retrained the decoder excluding the entire occipital lobe and yielded similar performance, indicating that the prediction is not driven by emotional schemas encoded in visual regions (Supplementary Results, Extended Data Fig. 3).

The interval between the picture presentation stage and the rating (button press) was fixed in the original disgust paradigm (Fig. 1b), which raises the possibility that the VIDS might partly decode the mapping of disgust onto motor responses. We therefore modified the disgust induction paradigm by including an additional jitter between the picture presentation and motor responding stage and further randomizing the order of rating 1 to rating 5 per trial (Supplementary Methods, Extended Data Fig. 4a). Applying the VIDS to the newly collected dataset (study 3, Supplementary Table 1) reveals that the disgust signature captures the experience of disgust rather than the mapping of disgust onto responses, such that the disgust signature yielded comparably high prediction-outcome correlations (within-subject *r*=0.87±0.02; overall prediction-outcome *r*=0.54) and could further predict different levels of disgust experience with high accuracy (details see Supplementary Results, Extended Data Fig. 4b).

To further determine the specificity of the VIDS to the experience of disgust (rather than anticipation or cognitive evaluation), we explored the VIDS reactivity over the stimulus presentation interval covering the pre-stimulus interval and onset of the stimulus in discovery (10×10-fold cross-validated) and validation cohorts. In line with the standard hemodynamic model, the VIDS response started approximately 4s following the picture onset and increased during 6-12s (Fig. 2d,e), suggesting that the VIDS captures disgust experience rather than expectation (pre-stimulus) or cognitive evaluation (response-reporting) of disgust (see also Supplementary Results, Extended Data Fig. 4c).

### The disgust signature predicts disgust across samples, paradigms and MRI systems

Accurate and population-level neurofunctional decoders require robust predictions across different paradigms, laboratories, and MRI systems. We therefore tested whether the VIDS could discriminate disgust versus neutral stimuli in a previously published independent study^54^ (study 4, Supplementary Table 1) that examined emotional reactivity to disgusting stimuli. The VIDS classified disgust versus neutral with high accuracy (92%, *P*<0.001, AUC=0.99, and Cohen’s *d*=1.46, Fig. 2f), indicating robust generalization to predict disgust experience across populations, experiments, and MRI systems.

### The disgust signature predicts the level of momentary disgust on the individual level

The conscious experience of disgust is a highly subjective and dynamic process^38,51^. A key question is thus to what extent the population-level VIDS can predict momentary variations in disgust on the individual level (on a trial-by-trial basis). To this end, single-trial models were employed to generate trial- and individual-specific activation maps (∼80 beta maps per subject in both cohorts). Next, we calculated the VIDS pattern expressions of these single-trial beta maps, which were finally correlated with the true ratings for each subject separately with statistical significance evaluated by prediction−outcome correlation (Pearson) for each subject. The results showed that the VIDS could significantly predict trial-by-trial disgust ratings in over 90% of the individuals in both, discovery (71 from 78, cross-validated) and validation (28 from 30) cohorts (Fig. 2g,h). Mean prediction−outcome correlations were 0.39±0.02 and 0.41±0.02 for discovery and validation cohorts, respectively. Our findings thus suggest that although disgust experience differs between individuals^38,51^, the VIDS could accurately and robustly predict the intensity of momentary disgust experience on the individual level across two independent populations.

### Subjective disgust is associated with and predicted by distributed neural systems

We next systematically determined which brain regions and systems contribute to the whole-brain disgust prediction. We first examined regions that made reproducible (reliable) contributions to the disgust prediction within the VIDS itself by applying a bootstrap test to identify regions with significant, consistent model weights (*q*<0.05, FDR corrected)^62^. Given that some brain features could contribute to controlling for noise in the data^55^ rather than the emotional process per se, we next transformed the population-level VIDS into ‘activation pattern’ (‘structure coefficient’, details see Methods). Results showed that a set of distributed subcortical and cortical brain systems exhibited significant model weights (Fig. 3a) and structure coefficients (Fig. 3b), including the bilateral amygdala, anterior and mid- and posterior-insular cortex, hippocampus, dorsal ACC, PAG, thalamus and putamen as well as the right mid-cingulate cortex, left posterior cingulate cortex, left inferior frontal gyrus, left superior temporal gyrus, right precentral gyrus, occipital and supplementary motor areas (Fig. 3c). Brain regions associated with and predictive of disgust experience on the individual level determined with convergent univariate and multivariate approaches identified a similar set of broadly distributed regions (Supplementary Methods, Supplementary Results and Extended Data Fig. 5). Overall, the analyses indicated that the conscious experience of disgust is represented in distributed subcortical and cortical systems involved in early detection and defensive responses towards potential threats (amygdala, PAG, thalamus, putamen)^8,18^, regions involved in interoceptive awareness, self-referential processing, and explicit emotional processing such as the anterior insula, dorsal ACC and frontal regions^29,30,44,63^, as well as regions that have been identified as core systems mediating the bitter taste reaction in animal models (posterior insula^6^, brainstem^5^). Negative associations with disgust were most consistently identified in superior frontal and middle temporal regions.

**Fig. 3.**
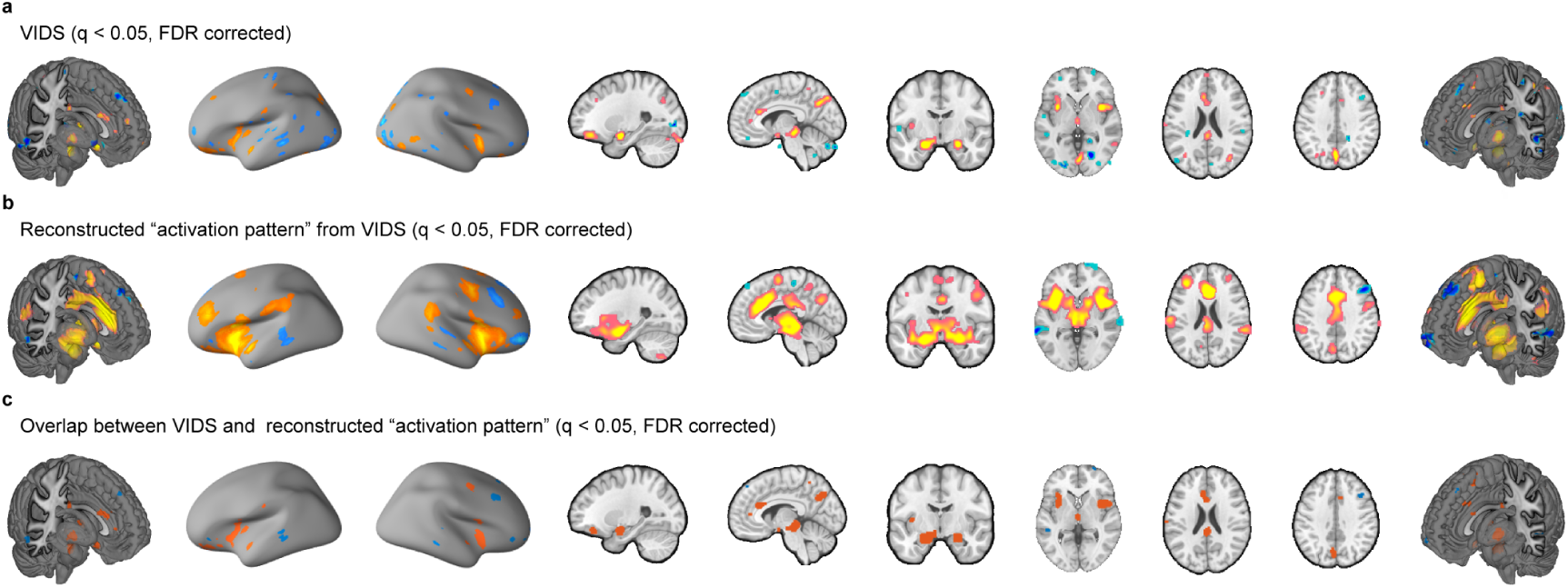
Subjective disgust experience is associated with and predicted by distributed brain regions. **a**, Thresholded VIDS. **b**, Thresholded transformed VIDS ‘activation pattern’. **c**, Conjunction between VIDS and transformed ‘activation pattern’. Images thresholded at *q*<0.05, FDR corrected. Hot color indicates positive weights (**a**) or associations (**b**) whereas cold color indicates negative weights (**a**) or associations (**b**).

### Spatial overlap of the neurofunctional disgust signature with the Brainnetome Atlas further confirmed a biological-plausible disgust model

To further evaluate the neurobiological plausibility and validity of the developed model, we calculated the spatial overlap percentage between the thresholded VIDS with the modified 279-region version of the Brainnnetome Atlas^64,65^ (see Methods). The top 15 atlas regions visualized in Fig. 4a included four amygdala sub-regions, five insula sub-regions, orbital gyrus, thalamus, rostral hippocampus, rostroventral cingulate gyrus as well as brainstem and midbrain regions involved in defensive responses and conscious emotional experience (i.e., PAG and substantia nigra^66^). Meta-analytic behavioural decoding utilizing the BrainMap database confirmed that the identified brain systems were primarily associated with negative emotional processes, in particular disgust showed the largest activation likelihood ratio (Fig. 4b,c). See Supplementary Results, Supplementary Table 2, and Extended Data Fig. 6 for the meta-analytic decoding results of the unthresholded VIDS pattern based on Neurosynth, which also supported the disgust signature as a biologically plausible model of disgust.

**Fig. 4.**
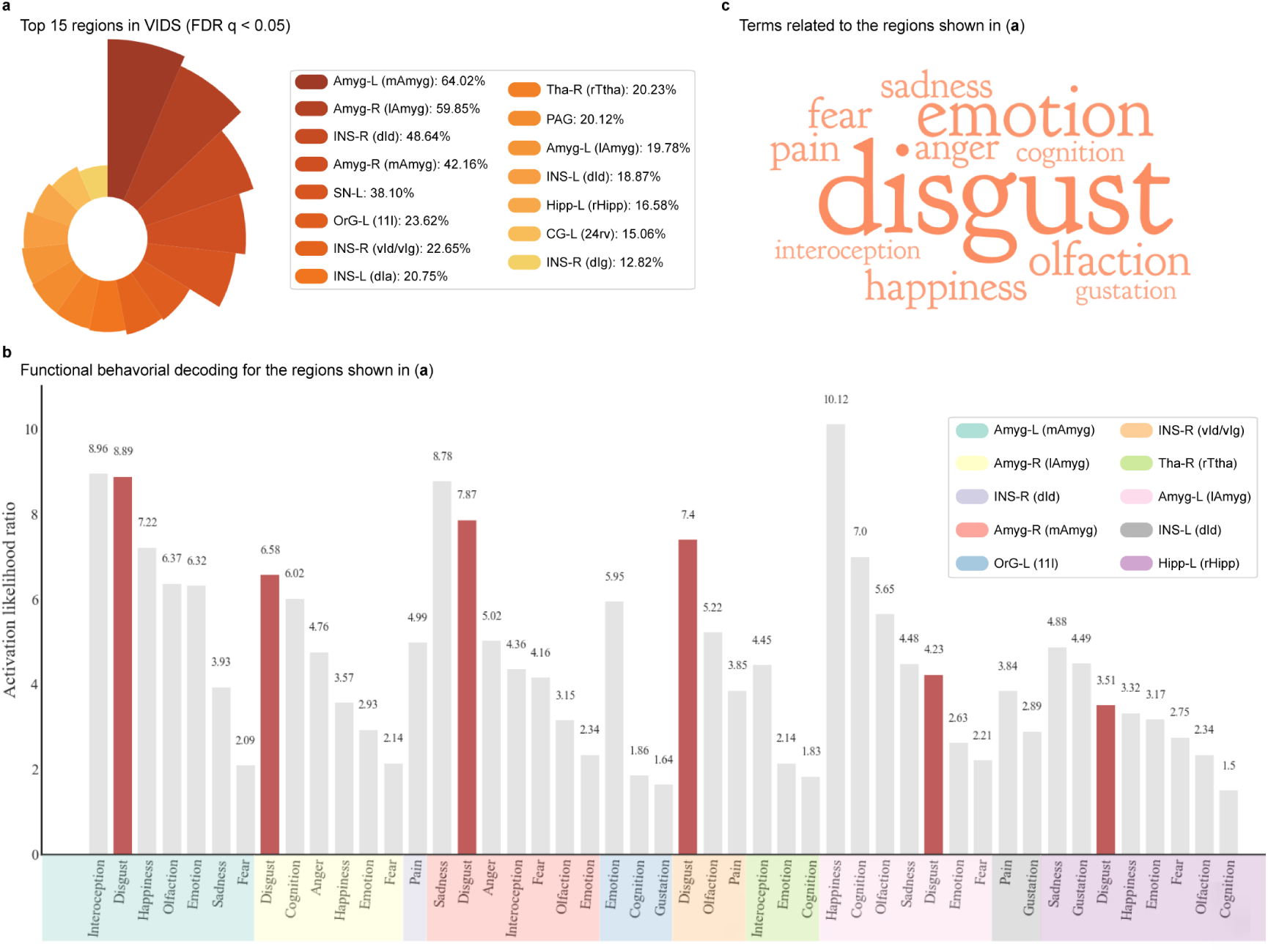
Neurobiological validity of the thresholded VIDS. **a**, The top 15 atlas regions overlapping with the thresholded VIDS (retaining positive values). **b**, Functional behavioural decoding for 10 exemplary regions for the thresholded VIDS (provided by the Brainnetome website). **c**, Meta-analytic behavioural decoding of the identified regions. Font size in the word cloud is proportional to the activation likelihood ratio related to each term provided by BrainMap, with larger activation likelihood ratio value depicted by larger font size.

### Alternative models to determine the contribution of isolated disgust-predictive systems: local searchlights, pre-defined regions, and networks show lower predictive accuracy compared to the VIDS

Given the continuing debate on the contribution of specific brain regions, such as insula^8,23,67,68^, amygdala^8,69^, and the DMN^70,71^, to subjective disgust experience, (1) both searchlight- and parcellation-based analyses were employed to determine local brain regions that were predictive of subjective disgust experience, and (2) models were trained on single brain region and network to examine to what extent these models could predict subjective disgust experience compare to the whole-brain VIDS. As shown in Fig. 5a,b, subjective disgust experience could be significantly predicted by activations in widely distributed regions (averaged across 10×10-fold cross-validation procedure). However, none of the local models predicted subjective disgust to the extent the VIDS did (see Extended Data Fig. 7a,b for predictions of models trained on discovery cohort on validation cohort).

**Fig. 5.**
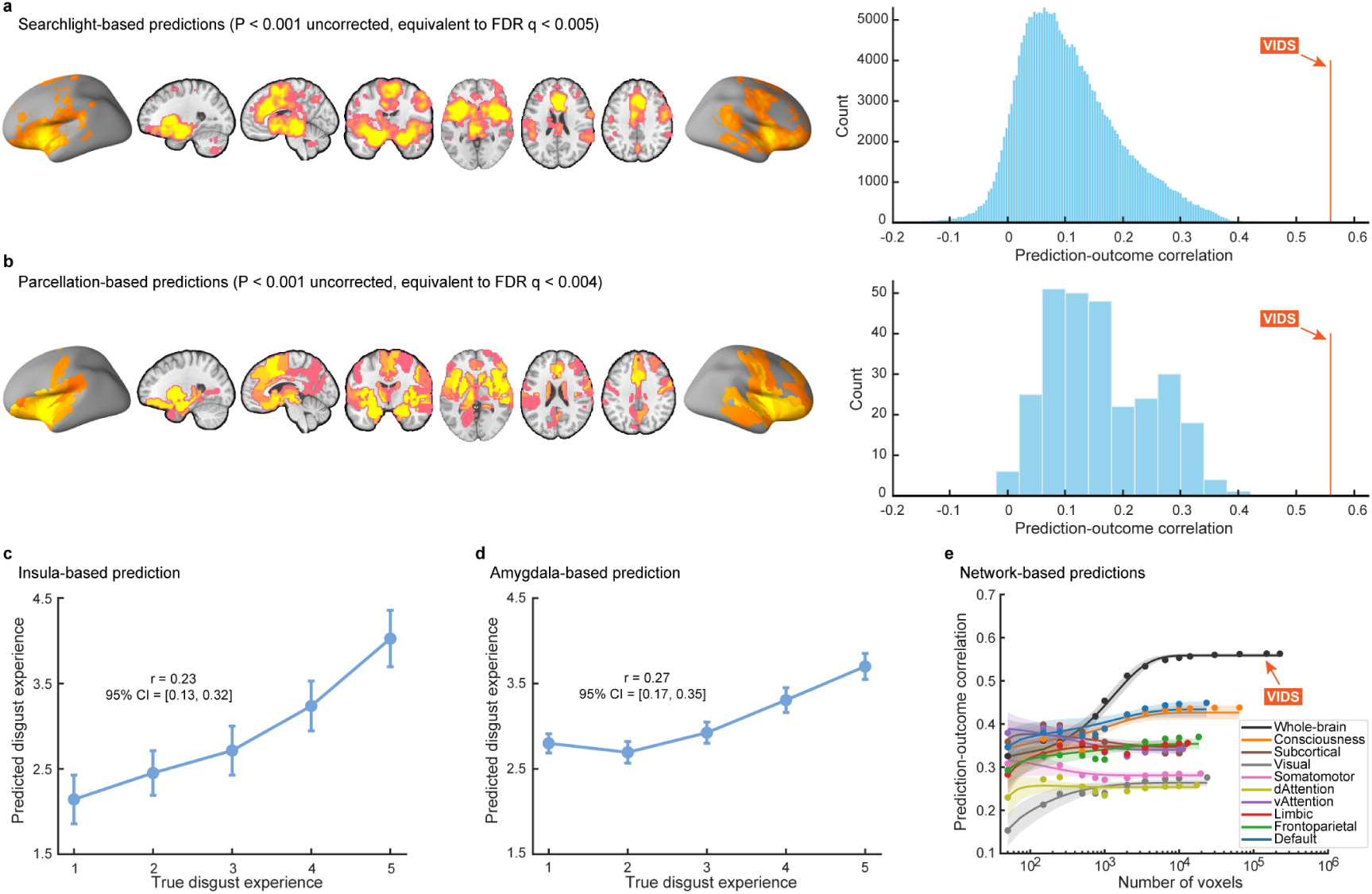
Local brain region and network predictions in the discovery cohort. **a,b**, Brain regions that significantly predict subjective disgust experience revealed by searchlight and parcellation-based analyses, respectively. Uncorrected *P* values equivalent to *q*<0.05 might be considered lenient, therefore brain regions that survived *P*<0.001 uncorrected (corresponding to *q*<0.005 and 0.004, FDR corrected, for searchlight- and parcellation-based predictions, respectively) are displayed. Histograms: cross-validated predictions (correlations) from local searchlights or parcellations. Orange lines indicate the prediction-outcome correlation from VIDS. **c,d**, Cross-validated predictions (mean±SE) from insula- and amygdala-based prediction analyses, respectively. Error bar indicates standard error of the mean; *r* indicates overall (between- and within-subjects; i.e., *n*=384 pairs) prediction-outcome Pearson correlation coefficient. **e**, Subjective experience of disgust is distributed across multiple systems. Model performance was evaluated as increasing numbers of voxels/features (x-axis) were selected to predict subjective disgust in different regions of interest including the entire brain (black), consciousness network (light orange), subcortical network (brown), or individual large-scale cerebral networks (other colored lines). The y-axis denotes the cross-validated prediction-outcome correlation. Colored dots indicate the mean correlation coefficients, solid lines indicate the mean parametric fit, and shaded regions indicate standard deviation.

We next re-trained predictive SVR models restricted to activations in (1) the bilateral insula; (2) the bilateral amygdala; (3) a consciousness network; (4) a subcortical network; (5) each of seven large-scale cerebral networks (see Methods). Results showed that the insula (prediction-outcome correlation *r*=0.23 and 0.38 for discovery (cross-validation) and validation cohorts, respectively), the amygdala (prediction-outcome correlation *r*=0.27 and 0.39 for discovery (cross-validation) and validation cohorts, respectively) as well as other brain networks (Fig. 5c,d,e, Extended Data Fig. 7c,d,e, and Supplementary Table 3) could, to some extent, predict subjective disgust experience. Nonetheless, although statistically significant (*P*s<0.001), the effect sizes in terms of prediction-outcome correlations (including searchlight- and parcellation-based predictions) were substantially smaller than those obtained from the VIDS, which used features spanning multiple brain systems. See also Supplementary Methods, Supplementary Results, and Extended Data Fig. 8 for separating a classically insula-associated process (pain empathy) from the representation of disgust experience.

To control for potential effects of the number of features/voxels in prediction analyses (i.e., the whole-brain model contains much more features), we randomly selected voxels (repeated 1,000 times) from a uniform distribution spanning the entire brain (black), consciousness network (light orange), subcortical (brown) or individual large-scale cerebral networks (averaged over 1,000 iterations)^18,40^. The asymptotic prediction when sampling from all brain systems as we did with the VIDS (black line in Fig. 5e and Extended Data Fig. 7e) was substantially higher than the asymptotic prediction within individual networks (colored lines in Fig. 5e and Extended Data Fig. 7e; see also Supplementary Table 3 for details). This analysis thus demonstrated that whole-brain models have much larger effect sizes than those using features from a single network. Moreover, model performance was optimized (i.e., reaching asymptote) when approximately 10,000 voxels were randomly sampled across the whole-brain, as long as voxels were drawn from multiple brain systems, further confirming that information about subjective disgust experience is contained in patterns of activity that span multiple systems. In addition, similar results were observed when applying the models trained on discovery cohort to the validation cohort (Extended Data Fig. 7e), indicating that models trained on approximately 10,000 randomly sampled voxels were robust enough (for similar findings, see ref.^18^). Together, converging lines of evidence from the above systematic analyses point to the fact that subjective disgust experience is encoded in distributed neural patterns that span multiple systems, adding to increasing evidence that emotions are represented in distributed brain systems rather than single brain regions or networks.

### The experience of disgust, fear, and non-specific negative affect have shared yet distinguishable neurofunctional representations

Disgust is inherently inter-related with general negative affect and functionally similar avoidance responses like fear^8^. To determine shared and separable neurofunctional representations, we conducted a series of analyses. First, we investigated spatial similarities between stable decoding maps and a set of ROIs as well as networks. Amygdala, anterior insula, basal ganglia, lPFC, and MCC contained stable predictive voxels in all three models, but VIDS showed larger contributions to the first four ROIs than VIFS and PINES while VIFS exhibited the largest contributions to MCC; furthermore, the remaining three ROIs (PAG, ACC, and thalamus) exhibited a degree of specificity for VIDS and VIFS (Fig. 6a). Although all networks showed stable predictive voxels across the three models, the relative contributions of each network to each model varied, such that the ventral attention, limbic, and consciousness networks contributed stronger to VIDS than to VIFS and PINES whereas the somatomotor network strongly contributed to PINES (Fig. 6b and Extended Data Fig. 9a). Second, we examined functional similarities between the VIDS and PINES^39^ (general negative emotion experience) and VIFS^18^ (fear), respectively. The results showed that VIDS was more specific to predict high versus low disgust as compared with PINES or VIFS, as reflected by effect sizes 1.56–2.76 and 2.09–2.61 times larger than those for PINES or VIFS in disgust discovery and validation cohorts, respectively (Fig. 6c). PINES was more sensitive to predict high versus low negative emotion with effect sizes 2.18–3.74 higher than those for VIDS or VIFS, and VIFS more accurately predicted high versus low fear with effect sizes 1.95–2.10 higher than those for PINES or VIDS (Fig. 6c). Particularly, additional analyses revealed that the three decoders were rather specific for their targeted emotion during discrimination at moderate versus low intensity (e.g., VIDS had effect sizes 1.84–27 and 2.31–11.53 times higher than those for PINES or VIFS in disgust discovery and validation cohorts, respectively (Fig. 6d). The above findings were further substantiated by the comparisons of the overall and within-subject prediction-outcome correlations of the three decoders across four datasets (Supplementary Table 4).

**Fig. 6.**
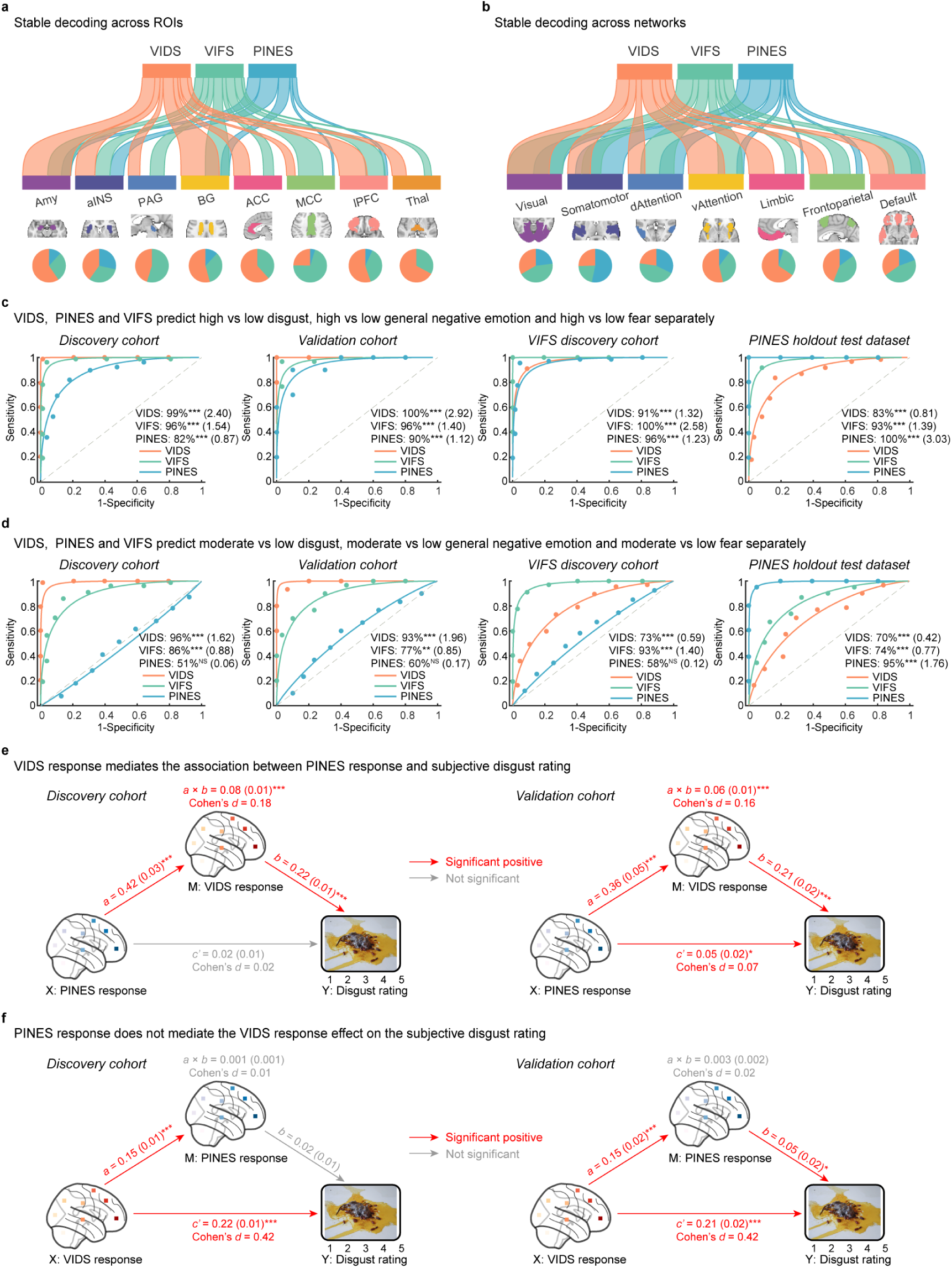
Comparing neurofunctional decoders for disgust, fear, and negative affect (VIDS, VIFS, PINES). **a,b**, River plots showing spatial similarity (cosine similarity) between stable decoding maps and selected brain systems. Ribbons are normalized by the max cosine similarity across all ROIs and networks. Stable decoding models were thresholded at FDR *q* < 0.05 and positive voxels were retained only for similarity calculation and explanation. Ribbon locations in relation to the boxes are arbitrary. Pie charts show relative contributions of each model to each ROI or network (i.e., percentage of voxels with highest cosine similarity for each map). **c,d**, VIDS, PINES, and VIFS more accurately (shown as forced-choice classification accuracy and Cohen’s *d*) predict the targeted emotion during discrimination at high versus low and moderate versus low intensities, respectively. **e**, VIDS response fully mediates the association between PINES response and subjective disgust ratings in the discovery cohort, and VIDS response partially mediates the PINES response - disgust rating association in the validation cohort. **f**, PINES response does not mediate the effect of VIDS response on disgust ratings, neither in the discovery cohort nor the validation cohort. **e,f**, The example picture from the DIRTI disgust images database^59^ (note that all pictures from the DIRTI are subject to creative commons license CC BY-NC 4.0); and the mediation analysis examines whether the observed covariance between the independent variable (X) and the dependent variable (Y) can be explained by the third variable (M, also mediator), details see Methods section. (**c,d**) ** *P*<0.01, *** *P*<0.001, NS not significant. (**e,f**) * *P*<0.05, *** *P*<0.001 (bootstrap tests with 10,000 samples; two-sided). aINS=anterior insula; BG=basal ganglia; MCC=midcingulate cortex; lPFC=lateral prefrontal cortex.

Finally, we employed multilevel mediation models, which examined whether the covariance between two variables (*X* and *Y*) can be explained by a third variable (*M*), to determine the neurofunctional relationship between the representations of subjective emotional experiences encoded in VIDS, PINES, and VIFS. While PINES could to some degree track disgust (discovery cohort: *r*=0.24; validation cohort: *r*=0.30), the VIDS response fully mediated the effect of PINES response on subjective disgust ratings in the discovery cohort, and in the validation cohort the VIDS response partially mediated the effect of PINES response on disgust ratings (Fig. 6e). In contrast, the PINES response failed to mediate the effect of VIDS response on disgust ratings (Fig. 6f). VIFS could predict subjective disgust (discovery cohort: *r*=0.35; validation cohort: *r*=0.32), and corresponding multilevel mediation models revealed that VIDS partially mediates the effect of VIFS response on disgust ratings and vice versa (Extended Data Fig. 9b,c); however, the first model (Extended Data Fig. 9b) had 3 times larger effect sizes than the second one (Extended Data Fig. 9c) in the discovery cohort and a similar trend was observed in the validation cohort, suggesting a stronger influence of VIDS in partially mediating the effects of VIFS response on disgust ratings. See Supplementary Results for further details.

Together, these findings underscore that neural representations of subjective experiences of negative emotions engage shared yet distinct representations which may shape distinguishable subjective experiences.

### The visual subjective disgust signature predicts gustatory stimulus induced (taste) disgust

From an evolutionary perspective, core disgust is considered to have evolved from the ancestral oral distaste response rooted in chemosensory rejection^1–3,11,14^. To test whether the visually-induced disgust signature generalizes to other modalities and whether the hypothesized link between distaste and subjective disgust is reflected on the brain level, the VIDS –as well as the PINES and VIFS – was applied to fMRI data acquired while participants tasted (as well as swallowed) either disgust or neutral liquids (study 7, *n*=30; Supplementary Methods and Supplementary Table 1). As shown in Fig. 7a, during the taste period, the VIDS could discriminate disgust taste from neutral taste with high accuracy (accuracy=80%, *P*=0.001, Cohen’s *d*=0.39) confirming that the VIDS generalizes to disgust across modalities, while neither the VIFS (accuracy=60%, *P*=0.36, Cohen’s *d*=0.26) nor the PINES (accuracy=53%, *P*=0.86, Cohen’s *d*=0.03) could significantly predict disgust taste versus neutral taste. Moreover, during the swallow period (Fig. 7b), the VIDS could significantly classify swallowing of disgust versus neutral liquids (accuracy=77%, *P*=0.005, Cohen’s *d*=0.68), while both the VIFS (accuracy=63%, *P*=0.20, Cohen’s *d*=0.43) and PINES (accuracy=50%, *P*=1, Cohen’s *d*=0.10) failed to predict the swallowing of disgust versus neutral liquids, further corroborating the specificity and sensitivity of VIDS for tracking disgust experience. Together, these results suggested that the visual disgust signature (i.e., VIDS) generalizes to the gustatory modality and exhibits a higher specificity and sensitivity compared to other signatures during both tasting and swallowing of disgust liquids.

**Fig. 7.**
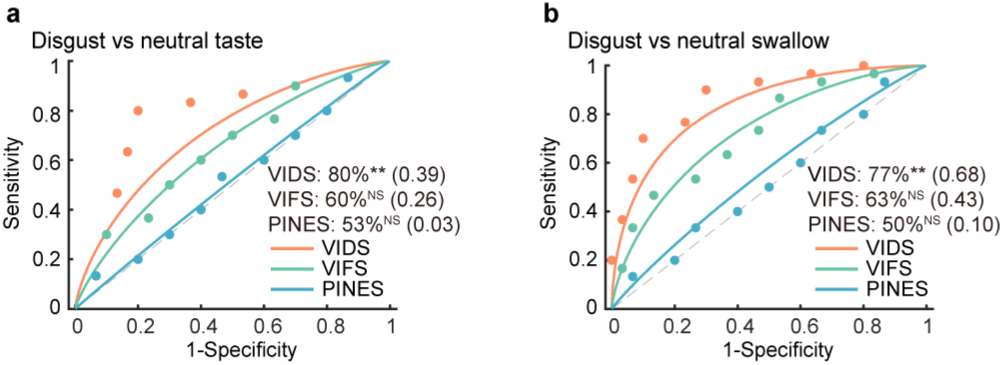
Validation tests of VIDS in the gustatory modality. **a**, (during the taste stage) VIDS could predict disgust versus neutral liquids (shown as forced-choice classification accuracy and Cohen’s d), while VIFS and PINES failed to predict disgust versus neutral liquids. **b**, (during the swallow stage) VIDS predicted disgust versus neutral liquids, however, neither PINES nor VIFS could predict disgust versus neutral liquids. ** *P*<0.01, NS not significant.

### The neural signature of subjective disgust experience captures sociomoral disgust in terms of neural reaction to unfairness

In everyday life, humans frequently refer to disgust experience in social (i.e., sociomoral) contexts and this has also entered pop-cultural contexts (e.g., ‘… I was ***disgusted*** by all the injustice, all the injustice…’ from Scream, Michael Jackson). However, debates in social and affective neurosciences continue on whether the evolutionary ‘core disgust’ that has evolved to avoid pathogens has been adopted towards social situations^8^, with some evidence showing that disgust experience evoked by moral transgressions (e.g., unfair offers) may be similar to the disgust experience elicited by core disgust stimuli (e.g., uncleanliness, pathogens)^3,72^, while other evidence suggested that unfair offers may evoke a general negative emotional response rather than disgust experience^53^. Here, we thus test whether the VIDS - which captures subjective disgust experience elicited by core disgust stimuli - could track sociomoral disgust. To this end, we performed exploratory predictions by applying the VIDS pattern along with PINES and VIFS to an independent fMRI dataset acquired while participants received a series of unfair offers in a social exchange (Ultimatum Game) task (study 8, *n*=43; Supplementary Methods and Supplementary Table 1). As shown in Fig. 8a,b, VIDS could predict high versus low unfairness with high accuracy (accuracy=74%, *P*=0.0019, Cohen’s *d*=0.73), and could also predict high versus moderate unfairness (accuracy=70%, *P*=0.0137, Cohen’s *d*=0.37). PINES could predict high versus low unfairness (accuracy=77%, *P*=0.0006, Cohen’s *d*=0.54) yet failed to predict high versus moderate unfairness (accuracy=56%, *P*=0.5424, Cohen’s *d*=0.20). VIFS did not predict either high versus low unfairness or high versus moderate unfairness. These results indicated that the neurofunctional reaction to unfair offers is highly similar to the emotional reaction towards core disgust stimuli.

**Fig. 8.**
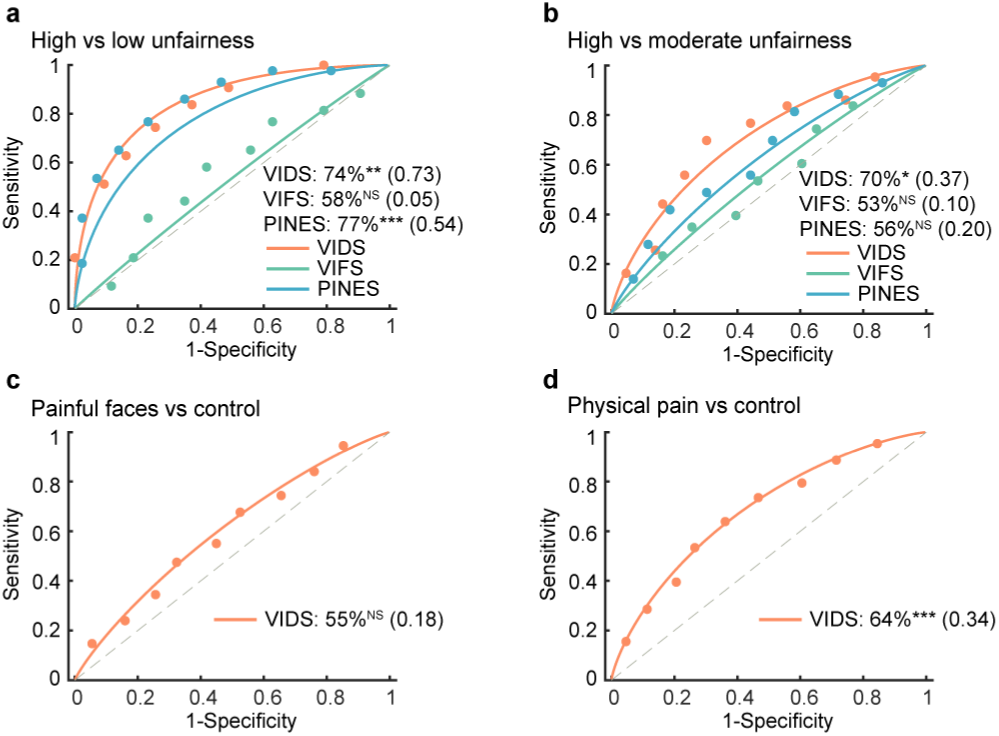
Validation tests of VIDS in social contexts. **a**, VIDS and PINES could predict high versus low unfairness (shown as forced-choice classification accuracy and Cohen’s d), while VIFS failed to predict high versus low unfairness. **b**, Neither PINES nor VIFS could predict high versus moderate unfairness, however, VIDS predicted high versus moderate unfairness. **c**, VIDS failed to predict painful faces versus respective control. **d**, VIDS could predict physical pain versus corresponding control. * *P*<0.05, ** *P*<0.01, *** *P*<0.001, NS not significant.

To further test whether VIDS simply captures negative emotional processing in high arousing social contexts, we applied the VIDS to another independent fMRI dataset (study 9, *n*=238; Supplementary Table 1) that acquired neural response during physical pain and painful facial stimuli as well as respective non-painful control stimuli. Results showed that the VIDS could not discriminate high arousing painful faces from respective control faces (with accuracy near chance-level, Fig. 8c). Although VIDS successfully distinguished physical pain versus corresponding control pictures (Fig. 8d), the effect size was substantially lower than that using VIDS to predict high versus low unfairness (Fig. 8a) (large effect sizes are commonly interpreted to indicate robust and replicable effects^73–75^); besides, the AUC of the VIDS for classifying high versus low unfairness (0.84) was larger than that for discriminating physical pain from corresponding control stimuli (0.68) (the AUC represents an important measure of classifier performance^46^). Combined, these findings indicated that VIDS is not sensitive to separate high arousing pain empathy from control stimuli, yet it is relatively specific for disgust experience in social contexts. For further analyses confirming that VIDS tracks unspecific arousal across negative and positive stimuli to a comparably low extent see also Supplementary Methods, Supplementary Results and Extended Data Fig. 10.

## Discussion

Recent perspectives propose a paradigm shift towards subjective and conscious emotional experiences in neuroscience^18,44,50,76,77^, yet neurobiological models that accurately describe the respective neural representations are scarce. Disgust originates in the hard-wired mammalian distaste reflex, but in humans its conscious and subjective emotional experience is considerably shaped by subjective appraisal and may extend towards sociomoral contexts and underlie some psychopathological dysregulations (e.g., obsessive-compulsive disorder). Utilizing machine-learning based neural decoding, we established and validated a comprehensive neurobiological model for subjective disgust experience (VIDS) that generalizes across samples and study contexts with regard to accurately predicting disgust in response to core disgust stimuli, gustatory distaste as well as unfair offers, suggesting that the experience of disgust may extend from the gustatory over the subjective experience and into the sociomoral domain (see also evolutionary models proposed in ref.^14^). Subjective disgust was encoded in distributed subcortical and cortical brain systems while no isolated region or network reached the predictive accuracy of the whole-brain model. In a series of analyses, we further demonstrated that subjective disgust experience showed shared yet separable neural representations with established predictive models for the subjective experience of non-specific negative affect or fear, such that all models engaged subcortical regions and regions implicated in emotional awareness and appraisal, whereas the neural signature of disgust robustly mediated the response of the other two models (PINES, VIFS) on subjective disgust ratings but not vice versa. Although the three neural signatures showed a certain extent of similarity in the range of intense emotional experiences (for a similar observation see also ref.^44^), the VIDS (as well as VIFS and PINES) were more sensitive to the target emotion in the moderate versus low intensity range. The VIDS moreover outperformed the other signatures in predicting disgust experience in response to both distaste and sociomoral disgust (unfair offers); while it tracked unspecific arousal in response to painful or positive pictures, the extent was comparably low. Together, the current study provides the first comprehensive neurofunctional model of subjective disgust, and the resulting VIDS can predict subjective disgust experience with high robustness and generalization.

Previous studies emphasized the role of single brain systems or networks as core modules for disgust, including the insula^7,8,22,23^ or the ventral attention network which may facilitate detection and attentional processing of core disgust stimuli^78^. Contrary to structure- and network-centric views, our findings indicate that subjective disgust experience requires concerted engagement of brain-wide distributed representations with comparably strong contributions of subcortical regions involved in rapid threat detection and avoidance responses (amygdala, PAG, thalamus, putamen)^8,18^ and cortical systems engaged in interoceptive awareness and emotional appraisal such as the anterior insula, dorsal ACC and lateral frontal regions^29,30,44,63^, as well as regions that have been identified as mediators of the bitter taste reaction in animal models (posterior insula^6^, brainstem^5^). However, neural activity in these isolated regions including systems such as insula or amygdala which have been typically implicated in disgust was only weakly associated with the subjective experience.

While no single network (e.g., visual network and ventral attention network) was critical for predicting subjective disgust, the consciousness network and DMN exhibited comparably strong contributions. According to two-system and higher-order consciousness models of emotion^63^, the consciousness network is vital for the generation and appraisal of conscious emotional states. In the context of constructionist theories of emotion, the DMN has been proposed to shape current affective experiences by drawing on autobiographical experiences and knowledge^70^ and to facilitate contextually sensitive escape decisions^79^. Consistent with previous predictive models for subjective emotional experiences^18^, our analyses also revealed that approximately 10,000 voxels that were randomly sampled across the whole-brain could lead to high predictive performance for subjective disgust. Together, our results dovetail with a growing body of research from both univariate^80^ and multivariate perspectives^18,39,42,43,52^ which demonstrated that capturing affective experience requires integration across multiple distributed neural systems. The distributed representation perspective aligns with appraisal^81^ and constructionist^51,80,82^ theories of emotions which propose that shared but also distinct distributed functional assemblies integrate to facilitate subjective emotional experiences.

In terms of the adaptive function, pathological dysregulations as well as subjective triggers and conscious experience level, disgust is - to a certain extent – distinguishable from related emotional domains like fear or negative affect (see e.g., also Research Domain Criteria, RDoC^83^). For robust fMRI-neuromarkers, a precise differentiation between subjective states is critical for model development and translation^44,46,48^. The VIDS exhibited partly overlapping yet also distinct neurofunctional representations compared to the PINES (non-specific negative affect)^39^ or the VIFS (fear experience)^18^. While prediction of all neuroaffective signatures was driven by contributions of distributed subcortical and cortical systems on the distributed signature level, the VIDS was more sensitive to predict subjective disgust than general negative affect or subjective fear. Moreover, the VIDS response mediated the association between PINES response and disgust ratings but not vice versa. Additionally, the correlation between VIFS response and disgust ratings was robustly mediated by the VIDS response. Together, these findings may underscore that the VIDS is predictive of subjective disgust experience rather than predicting unspecific salience or arousal or non-specific aversive emotional states and that shared yet also separable neurofunctional representations shape subjective emotional experiences.

According to Rozin et al., the distaste mechanism directly activates the disgust output system without a disgust evaluation stage^1^, reflecting a reflexive hard-wired response to the chemosensory properties of gustatory stimuli. Mice do exhibit defensive responses (e.g., retching-like behaviour)^5^ and distinct facial expressions^6^ after ingestion of potentially poisonous food, non-human primates may expel unpleasant tasting food^10^, and human neonates exhibit a characteristic facial grimace towards bitter tastes^84^. The present study provides evidence for a link between the distaste response and the subjective core disgust experience by showing that the predictive potential of the signature for visually-induced disgust generalizes to the neural representation of the gustatory distaste response (not only during the taste but also the swallow of disgust liquids) and provides supporting evidence for the proposed origin of disgust in oral distaste. Interestingly, meta-analytic functional decoding based on the Neurosynth repository not only revealed an association with affective states, including disgust (e.g., disgust, feelings) but also an association with several gustation-related terms (e.g., eating, food, taste; Extended Data Fig. 6).

We next utilized the disgust predictive model to determine whether disgust experience extends into the sociomoral domain^3,53,72^, in particular whether exposure to unfairness activates the VIDS. Evolutionary and disgust appraisal models have long hypothesized an association^1,14^, but the lack of a precise neuromarker for subjective disgust precluded a direct test of the hypothesis. Chapman et al. thus emphasized that the MVPA approach is suitable to examine the extent of core disgust appraisal being involved in sociomoral disgust as it provides evidence for the activation of shared appraisal while it is concomitantly critical to decode other aversive states associated with withdrawal (e.g., fear)^14^. Our results revealed that VIDS could predict different levels of unfairness with high accuracy (high, moderate, low unfairness) while PINES could differentiate high unfairness from low unfairness yet failed to predict high versus moderate unfairness, and VIFS did not react to unfairness. These findings largely converged with the observations from Chapman et al.^3^. The authors showed that participants judged their experience as most similar to disgust when they received unfair offers, while other emotions like fear did not change with unfair offers. More importantly, however, the authors noted that the emotional response to unfairness was not characterized by disgust alone; rather, it is a complex blend of multiple negative emotions, and anger and sadness may also play a role, albeit to a lesser degree than disgust^3^. Indeed, through item analysis, Chang et al. found that the PINES response was strongly associated with normative ratings of multiple discrete negative emotions (e.g., disgust, *r*=0.94; anger, *r*=0.94; sadness, *r*=0.92)^39^, which explains why the PINES also captured unfairness to a certain extent as revealed in the present study. Unfair behaviour could thus induce a broad range of negative emotional responses in the observer. Therefore, we constructed a route from unfairness to sociomoral output via ‘other negative affect evaluation’ in Fig. 1a. Notably, a previous study employing MVPA also emphasized the role of general negative affect in response to unfairness^53^ by demonstrating shared regional neural representations in the insular-mid ACC system for disgust and unfairness, a finding that is partly mirrored in the observation that the PINES could differentiate high versus low (yet not high versus moderate) unfairness (Fig. 8a). Other recent findings suggest that moral cognition interacts in a privileged manner with a neural representation of disgust and carefully controlled for aversiveness (i.e., pain)^85^. The same study further underscores the role of functional interactions between regions as the common neural link between moral cognition and core disgust, such that enhanced coupling between separate subsystems (e.g., PCC versus left ventral anterior insula) may link the neurofunctional pathway underlying disgust stimulation and moral appropriateness ratings of ethical dilemmas^85^. While the different paradigms used in the studies (directly experienced unfairness in the Ultimatum game in the present study versus a cognitive evaluation of ethical dilemmas in ref.^85^) may have contributed to the differences in determining shared neural activity patterns between disgust and sociomoral processes, future studies may include data acquired during different social cognitive and disgust processes and simultaneously account for shared and separable distributed activity and connectivity features (for a synergistic decoding approach see also ref.^86^). Altogether, our findings extend the initially proposed disgust appraisal model depicted in Fig. 1a, which combined several possibilities raised in previous disgust appraisal models^1,14^. Rozin et al. once mentioned that ‘…only if evidence is found for a route from unfairness to the disgust evaluation system can it be concluded that disgust at unfairness is “the same” as disgust that is elicited through the core route (such as in response to cockroaches) …’^1^. We here demonstrate initial evidence for a route between the unfairness and the subjective disgust evaluation system. From the evolutionary perspective, these findings support the notion that disgust may have been preadapted to avoid harm and regulate behaviour in social contexts (see also adaptationist theory by Tybur et al.^11,12^).

From a biomarker perspective, it is imperative that a neuroaffective signature captures the respective mental process across variations of experimental contexts^40^. The VIDS generalized across cohorts, core and sociomoral disgust (as well as gustatory distaste) paradigms, MRI systems and predicted disgust with considerably higher effect sizes than reactivity to high arousing positive or high arousing negative stimuli or strongly aversive vicarious pain stimuli (pain infliction stimuli). See Supplementary Results for further discussions on the arousal issue. Since the insula plays an important role in encoding pain empathy and disgust^49,53,87^, we further tested a physical pain empathy pattern^49^ on discovery and validation cohorts, yet the performance was substantially inferior to VIDS (see Extended Data Fig. 8). Together with the in-depth comparison of the predictive and topographical features of the fear, non-specific negative affect, and disgust signatures, we demonstrate that the VIDS is a comparably robust, sensitive, and specific brain marker to track and differentiate subjective disgust experience across contexts.

The present study has the following limitations. While we observed partly overlapping systems between the subjective experience representations of disgust and fear, the extent to which these common representations represent hard-wired physiological defensive responses, autonomous reactivity, or general emotional arousal remains unclear. Furthermore, this study performed explanatory predictions by testing the developed VIDS, together with VIFS and PINES, in the sociomoral domain; however, we could not provide a direct prediction of the degree of other negative emotions like anger engaged throughout this process (for the role of anger in sociomoral disgust, see refs.^14,88^). While the current study supports models that posit a strong link between core and sociomoral disgust, we focused on unfairness within a highly specific domain because aligning with fairness norms is deemed as a central part of human morality and sociality^3,89,90^. Nevertheless, moral disgust experience may extend to a range of social situations and future studies may determine whether neurobiological models based on distributed activation patterns or coupling between regions^85^ generalize across domains. Combining fMRI with MVPA enables us to detect and characterize fine-grained activity patterns with considerably higher precision compared to the conventional univariate approach, and can extend our understanding of how the brain engages in experimental tasks^56,91^. However, the fMRI/MVPA combination still requires cautious interpretation given that MVPA cannot compensate for the inherent limitations of fMRI which indirectly probes neuronal activity, and distributed activation pattern cannot be viewed as reflecting the specific computational process of the neurons within a voxel^40,45,47^. Future studies should aim at providing more precise measurements of how fMRI aligns with detailed neuronal recordings at the microscopic scale (see also discussion in ref.^91^).

In conclusion, the present study developed a whole-brain neural signature for subjective disgust experience. This visually-induced disgust pattern was validated and generalized across participants, paradigms, study contexts, and MRI systems. We showed that the neural basis of subjective disgust is encoded in multiple distributed (large-scale) brain systems rather than isolated brain regions. The robustness and specificity of the resulting VIDS were further tested with general negative emotion experience, subjective fear experience, gustatory modality (distaste), and social contexts (e.g., pain empathy and sociomoral disgust). This study adds to the understanding of the neurobiological underpinnings of subjective disgust experience. Disgust dysregulations play a role in a number of mental disorders and the resulting subjective disgust marker can be potentially useful to examine novel treatments (for initial applications of MVPA-based decoders in treatment and intervention evaluation, see refs.^21,92^).

## Methods

### Participants in the discovery cohort (Study 1)

Discovery cohort (labeled as study 1) included eighty participants recruited from the University of Electronic Science and Technology of China (UESTC). All participants were right-handed, reported having normal or corrected to normal vision, had no history of mental or physical disorders, no MRI contraindications, and were free of current or regular substance or medication use. Data from two participants (one female) were excluded due to excessive head movement (>3mm) during fMRI scanning, therefore, leading to a final sample of *n*=78 participants (44 females; mean±SD age=22.10±2.68 years). Informed consent was obtained before the experiment, the experimental protocol was approved by the UESTC Ethics Board and in line with the latest revision of the Declaration of Helsinki. Participants were reimbursed 110 RMB.

### Participants in the validation cohort (Study 2)

In order to validate the performance of the disgust decoder (VIDS) developed using the discovery cohort (i.e., study 1), we next collected an independent dataset (validation cohort, study 2) using the identical paradigm as in the discovery cohort. The data acquisition plan and analyses were pre-registered before the start of the validation experiment (https://osf.io/y4s78). Study 2 included thirty-one healthy participants recruited from the UESTC. Enrollment criteria were identical to study 1. Data from one male participant were excluded due to excessive head movement (>3mm) during fMRI scanning, leading to a final sample of *n*=30 participants (16 females; mean±SD age=21.13±2.18 years). Informed consent was obtained before the experiment, the experimental protocol was approved by the UESTC Ethics Board and in line with the latest revision of the Declaration of Helsinki. Participants were reimbursed 110 RMB.

### Stimuli and paradigm used in the discovery cohort (Study 1)

The disgust rating task incorporated 80 pictorial stimuli from the Disgust-related Images (DIRTI) database^59^ (67 pictures), Nencki Affective Picture System (NAPS)^93^ (5 pictures), and International Affective Picture System (IAPS)^94^ (8 pictures), distributing over 4 runs (with 20 stimuli per run). Of note, to avoid specificity driven by different categories of disgust, we drew a balanced sample from animals, humans, and scenes. Stimuli were presented using the E-Prime stimulus presentation software (Version 2.0; Psychology Software Tools, Sharpsburg, PA). Participants were instructed to pay attention to the pictures and naturally experience the induced emotion. Each trial consisted of a 6s presentation of the picture followed by a 2s fixation-cross separating the stimuli from the rating period. Participants then had 4s to report the level of disgust they experienced for the stimuli using a 5-point Likert scale where 1 indicated neutral/slightest disgust and 5 indicated most strongly disgust. Finally, there was a jittered fixation-cross epoch (5, 6, or 7s) before the presentation of the next picture (Fig. 1b). All participants reported ‘1-4’ in their responses while 6 out of 78 participants did not use rating ‘5’.

### Stimuli and paradigm used in the validation cohort (Study 2)

Stimuli and paradigm used in the validation cohort were identical to the discovery cohort. All participants reported ‘1-4’ in their responses while 1 out of 30 participants did not use rating ‘5’.

### MRI data acquisition for discovery and validation cohorts

MRI data were acquired on a 3T system (GE MR750, General Electric Medical System, Milwaukee, WI, USA). Structural images were collected using a high-resolution T1 spoiled gradient recall (SPGR) sequence (repetition time=6ms, echo time=2ms, flip angle=9°, field of view=256×256mm, 1×1×1mm voxels, 256×256 acquisition matrix, 156 slices, 1mm slice thickness) and were used for improving spatial normalization and excluding participants with apparent brain pathologies. Functional images were acquired with a T2*-weighted echo planar imaging (EPI) sequence (repetition time=2000ms, echo time=30ms, flip angle=90°, field of view=240×240mm, 3.75×3.75×4mm voxels, 64×64 resolution, 39 slices, 3mm slice thickness, 1mm gap).

### fMRI data preprocessing in discovery and validation cohorts

In line with our previous studies^18,49^, all fMRI data were preprocessed and analyzed using SPM12 (Statistical Parametric Mapping, https://www.fil.ion.ucl.ac.uk/spm/software/spm12/). The first five volumes of the fMRI data in each run were removed to allow for T1 equilibration. Briefly, the preprocessing steps included: correction for differences in slice timing, realignment of head motion, unwarp to correct for magnetic field inhomogeneity, followed by the tissue segmentation of high-resolution anatomical image (into gray matter, white matter, cerebrospinal fluid, bone, fat and air), co-registration of the skull-stripped and bias-corrected structural image to the functional images, normalization of functional images to the standard Montreal Neurological Institute (MNI) space (interpolated to 2×2×2mm^3^ voxels) and spatial smoothing with an 8-mm full width at half maximum (FWHM) Gaussian kernel. In line with earlier studies (e.g., refs.^18,49,60,95,96^), we used smoothed data (8-mm FWHM Gaussian kernel) for MVPA as previous studies indicated that this smoothing level could improve inter-subject functional alignment without impairing the sensitivity to mesoscopic activity patterns that are consistent across participants^97,98^. Before first-level analysis, global motion outliers in each run were identified according to any of the following criteria: (a) signal intensity exceeding three standard deviations from the global mean or (b) signal intensity and Mahalanobis distances exceeding ten mean absolute deviations (https://github.com/canlab/CanlabCore). Each time point identified as outliers was later included as a separate nuisance covariate in the first-level model.

### First-level fMRI analysis in discovery and validation cohorts

We designed two separate participant-level univariate general linear models (GLM). Specifically, the first GLM model used to create beta images for the following prediction analyses included five separate regressors time-logged to the presentations of pictures for each rating (i.e., 1-5), enabling us to model brain activity in response to each disgust level separately. The model also included one boxcar regressor indicating the 4s rating period aiming to model effects correlated with motor activity. The fixation cross epoch was used as an implicit baseline. The second GLM model employed a parametric modelling approach using subjective disgust ratings encompassing two regressors of interest, with one modeling the picture presentation period and the other modeling the subjective disgust rating period. The self-reported disgust ratings (i.e., 1-5) for each picture were included in the statistical model as a parametric modulator for the presentation period. The model thus allowed to determine brain regions where neural activity changed as a function of disgust ratings.

Task regressors were convolved with the canonical hemodynamic response function and a high-pass filter of 128s was applied. The four runs of the fMRI task were concatenated for each participant (via SPM spm_fmri_concatenate.m function). Nuisance variables encompassed: (a) ‘dummy’ regressors representing each run (intercept for each run); (b) the six estimated head movement parameters (*X*, *Y*, *Z*, roll, yaw, and pitch), their squares, their derivatives and squared derivatives for each run (24 columns in total); (c) vectors indicating motion outlier time points.

### Multivariate pattern analysis

We used whole-brain multivariate machine-learning pattern analysis^18,39,43,62^ to determine a pattern of weighted brain activity that best predicted the self-reported disgust ratings. Here, we used the support vector regression (SVR) algorithm (linear kernel with *C*=1) implemented in the Spider toolbox (http://people.kyb.tuebingen.mpg.de/spider). Whole-brain data masked with a gray matter mask from the discovery cohort was used. Features (i.e., predictors) constituted 384 beta maps (one per rating for each participant, i.e., 72×5 + 6×4), aggregated into an images × voxels matrix (stacked across participants), and the outcome variable included subjective disgust ratings across participants. To test the performance of the disgust-predictive pattern and to rule out the possibility of model overfitting^21,39,99^, we used a rigorous 10×10-fold cross-validation procedure on the discovery cohort during which all participants were randomly assigned to 10 subsamples of 7 or 8 participants using MATLAB’s cvpartition function. The optimal hyperplane was computed based on the multivariate pattern of 70 or 71 participants (training set) and evaluated by the excluded 8 or 7 participants (test set). This procedure was repeated ten times with each subsample being the testing set once. To avoid a potential bias of training-test splits, the cross-validation procedures throughout the study were repeated ten times by producing different splits in each repetition and the resultant prediction performance were averaged to produce a convergent estimation^18,49,100^.

To further evaluate the performance of the disgust-predictive pattern (trained on the whole discovery cohort), we assessed overall (between- and within-subjects; 384 pairs in total) and within-subject (5 or 4 pairs per subject) Pearson correlations (*r*) between the cross-validated predictions and the actual ratings to indicate the effect sizes and the RMSE to illustrate overall prediction error (for similar model evaluation metrics, see ref.^43^). Furthermore, we computed the classification accuracies of the VIDS between each successive pair of disgust rating levels (rating 2 versus 1, rating 3 versus 2, rating 4 versus 3, and rating 5 versus 4) as well as between low, moderate and high disgust rating levels (i.e., high versus low, high versus moderate, and moderate versus low) from receiver operating characteristic curves using forced-choice classification, where signature responses were compared for two conditions tested within the same participant (the higher was chosen as more disgust) and is therefore ‘threshold free’. Two-sided binomial tests were used to test whether the classification accuracies were higher than chance-level (50%).

Additionally, to test how well the VIDS predicted disgust on datasets other than the discovery cohort (and to rule out the possibility of overfitting^99^), we applied the VIDS to multiple separate test datasets, e.g., the validation cohort, the dataset from the modified disgust induction task (study 3, Supplementary Methods), the generalization cohort (see ‘Generalization cohort’ section below), gustatory disgust dataset (see ‘Validation in the gustatory context’ section below), and Ultimatum Game dataset (see ‘Validation in social contexts’ section below) to obtain a pattern response for each map (that is, the dot-product of each vectorized univariate GLM-derived fMRI activation map with the VIDS weight map).

### Generalization cohort

A previous study measured disgust experience and regulation which included disgust and neutral stimuli (study 4^54^). In line with the aim of the current study, we focused on the disgust experience condition. Briefly, 26 participants (10 females; mean±SD age=21.73±1.69 years) performed a disgust experience task where they were required to pay attention to the pictures shown on the screen. There were 10 blocks (5 disgust and 5 neutral blocks), and each block encompassed 3 consecutive pictures (2s each). After each block, the participants were required to rate how negative they felt. Given that the ratings were averaged across several pictures, we focused on the classification of disgust versus neutral to increase statistical power. See Supplementary Methods for the MRI acquisition details.

### Within-subject trial-wise prediction

Here we tested whether the VIDS could predict individual trial-by-trial subjective disgust. To this end, we performed a single-trial analysis^18,101,102^, which was achieved by constructing a GLM design matrix with separate regressors for each stimulus. Each task regressor was convolved with the canonical hemodynamic response function. Nuisance regressors and high-pass filter were identical to the above GLM analyses. When using a single-trial analysis, it is worth noting that the regressor (i.e., beta estimates) for a given trial could be strongly contaminated by artifacts (e.g., participant’s head movement, scanner pulse artifacts, etc.) that co-occur during that trial. Given this, for each subject, variance inflation factors (a measure of design-induced uncertainty due to collinearity with nuisance regressors) were calculated for each regressor, based on whether it can be predicted by a combination of the others. And any trials with variance inflation factors over 3 were excluded from subsequent analyses (overall ∼3.7% of trials in the discovery cohort as well as ∼3.4% of trials in the validation cohort were excluded). Next, these single-trial beta maps were used in VIDS pattern expression analysis (i.e., the dot-product of vectorized activation images with the VIDS weights). For subjects in the discovery cohort, a 10×10-fold cross-validation procedure was employed to obtain the VIDS response of each single-trial beta map for each subject.

### Determining brain regions associated with and predictive of subjective disgust

A series of analyses were employed to identify neural substrates underlying subjective experience of disgust. Firstly, we conducted bootstrap tests to determine brain areas that made stable contributions to the prediction across participants, where we constructed 10,000 samples (with replacement) from the discovery cohort, repeated the prediction process with each bootstrap sample, and evaluated *z*-scores and two-tailed uncorrected *P* values at each voxel based on the mean and standard deviation of the bootstrap distribution, on the population-level disgust-predictive pattern (i.e., the VIDS). After correction for multiple comparisons (FDR *q*<0.05), those regions with significant and consistent model weights were obtained^62^. Secondly, as in previous studies that computed model encoding maps^18,43^, we transformed the population-level pattern (i.e., VIDS, also backward model) into ‘activation pattern’ (forward model), which maps each voxel to the response (fitted values) in the multivariate model^55^. The reconstructed ‘activation pattern’ is known as ‘structure coefficient’ in the statistical literature^103,104^. And the one-sample *t* test, thresholded at FDR *q*<0.05 corrected for multiple comparisons, identified reliable disgust predictive and associative brain regions of the VIDS. Core regions that were viewed as the most consistently correlated with and predictive of the target outcome (i.e., subjective disgust) were defined as having voxels that were consistently observed across backward (that is, model predictive weights) and forward (that is, model encoding voxels in which the prediction correlates with fMRI activation) models (for the methodology detail of determining regions involved in the subjective experience of disgust on the individual level, see Supplementary Methods).

Next, to independently evaluate the neurobiological validity of the developed pattern, we adopted two approaches. First, we used the thresholded VIDS (FDR *q*<0.05, retaining positive values) from the bootstrap test and examined its overlap percentages with the modified 279-region version of the Brainnetome Atlas included additional brainstem, midbrain, and cerebellum regions^64,65^. We then displayed the top 15 regions. Moreover, to further decode the functions correlated with these regions in the thresholded VIDS, functional characterizations of ten exemplary regions as provided by the Brainnetome website (https://atlas.brainnetome.org/bnatlas.html) were reported through probabilistic maps indicating the behavioural domain based on meta data labels of the BrainMap database (http://www.brainmap.org/taxonomy/)^64^. Second, to complement the BrainMap functional decoding, we further performed an exploratory decoding analysis of the disgust pattern using data derived from Neurosynth and NiMARE^105,106^ which contain a large pool of automatically generated meta-analytic activation maps across a multitude of terms/topics. This approach enables us to discuss our results in relation to these terms/topics, without relying on acquiring data from a wide range of functional neuroimaging tasks in the same cohort. Consistent with the recommended input specifications as well as recent publications^62,107–109^, we subjected the unthresholded VIDS pattern for the meta-analytic decoding. Here, we capitalized on the BrainStat toolbox (https://github.com/MICA-MNI/BrainStat)^110^ to run the analysis. For every meta-analytic map in the database, a voxel-wise Pearson product-moment correlation between the meta-analytic map and input map was calculated. We only displayed the top 100 most correlated functional terms/topics for clarity. Additionally, we excluded topics such as dorsolateral prefrontal, anterior cingulate, etc.

Furthermore, we tested whether disgust processing could be reducible to activations in a single brain region (e.g., insula, amygdala) or network (e.g., the DMN). To examine this hypothesis, we performed whole-brain searchlight (three-voxel radius spheres) – and parcellation (279 regions^64,65^) – based analyses to identify local regions predictive of disgust and compared model performances of local regions with the whole-brain model (i.e., the VIDS). Additionally, we compared the prediction performances of insula and amygdala (based on the 279-region version of the Brainnetome Atlas^64,65^) as well as large-scale networks to the whole-brain model. The networks of interest comprised of seven large-scale cerebral networks^111^, a subcortical network (including the striatum, thalamus, hippocampus, and amygdala), and a cortical ‘consciousness network’^63^. To reduce potential biases arising from different atlases, we continued to use the modified 279-region version of the Brainnnetome Atlas (which also combined Yeo’s seven networks) to extract the nine networks. For these analyses, we trained and tested a model for each searchlight sphere, parcellation, brain region, or network separately using the discovery data (10×10-fold cross-validated).

### Comparing the performance of VIDS with the PINES and the VIFS

Previous studies have developed and evaluated whole-brain emotional decoders for general negative emotion experience (picture-induced negative emotion signature, PINES^39^) and subjective fear experience (visually-induced fear signature, VIFS^18^). To compare the performance of VIDS with the PINES and the VIFS, we applied the three decoders to the discovery, validation, PINES holdout test^39^ (Study 5, *n*=61, see Supplementary Table 1 for details) and VIFS discovery^18^ (study 6, *n*=67, Supplementary Table 1) cohorts and assessed the overall as well as within-subject prediction−outcome correlations between the pattern expressions and the true ratings. Two-alternative forced-choice classification accuracies between the separate disgust intensity levels (and general negative emotion as well as fear) based on the pattern expressions were further calculated.

### Spatial similarity between stable decoding maps and a priori regions of interest as well as networks of interest

River plots were created to illustrate spatial similarity between stable decoding maps derived from bootstrap tests and a prior regions of interest previously documented as regions linking to disgust, fear, and negative affect processes^8,18,39,43^. We further depicted spatial similarity between stable decoding maps and seven large-scale cerebral networks, the subcortical and consciousness networks. In line with a recent study^43^, spatial similarity was computed as cosine similarity between the ROI or network and the thresholded VIDS, VIFS, and PINES (FDR *q*<0.05, retaining positive values) from bootstrap tests.

### Multilevel two-path mediation analysis

To explore the relationship between VIDS response, disgust rating, and PINES response, multilevel two-path mediation analyses were performed using the Mediation Toolbox, available via https://github.com/canlab/MediationToolbox^18,112^. Briefly, the mediation analysis examines whether the observed covariance between the independent/predictor variable (*X*) and the dependent/outcome variable (*Y*) can be explained by the third variable (*M*, also mediator). The predictor-mediator relation, mediator-outcome relation, and predictor-outcome relation before and after controlling for the mediator are characterized by paths *a*, *b*, *c,* and *c’*, respectively. Specifically, the total effect of the predictor on the outcome (path *c*) is the sum of direct/non-mediation effect (path *c’*) and indirect/mediation effect (the product of the path coefficients of path *a* and path *b*, i.e., *a*×*b*). Significant mediation effect is obtained when *a*, *b,* and *a*×*b* are all significant. Furthermore, when *c’* is significant, *M* (i.e., the mediator) is considered to have a partial mediation effect; otherwise, *M* plays a full mediation role. In this study, we constructed two multilevel mediation analyses: (1) the trial-by-trial VIDS responses were entered as predictors (*X*), disgust ratings were entered as outcomes (*Y*), and the trial-by-trial PINES responses were entered as mediators (*M*); (2) the trial-by-trial PINES responses were entered as predictors (*X*), disgust ratings were entered as outcomes (*Y*), and the trial-by-trial VIDS responses were entered as mediators (*M*). To do this, the VIDS and PINES responses were calculated by dot-product of the single-trial beta maps (trials with variance inflation factors over 3 were excluded) with the VIDS and PINES patterns, respectively, for each participant. Bootstrap tests with 10,000 iterations were used to assess the statistical significance of mediation effects. If the bootstrapped 95% CI does not include zero, the effect will be considered to be significant (*P*<0.05). Of note, we also tested whether (1) VIFS response mediated the relationship between VIDS response and disgust ratings; and (2) VIDS response mediated the relationship between VIFS response and disgust ratings.

### Validation in the gustatory context

To test whether the VIDS – developed on visual stimuli – can generalize to the gustatory modality, we designed and implemented a new fMRI experiment (study 7, *n*=30; Supplementary Methods and Supplementary Table 1) employing gustatory stimuli (design based on previous similar studies^3,53,113–115^). Next, we applied the VIDS –as well as the VIFS and PINES - to the gustatory fMRI data. Specifically, we calculated forced-choice classification accuracies between disgust and neutral taste (as well as between swallowing of disgust and neutral liquids) based on the pattern responses.

### Validation in social contexts

First, to examine whether the VIDS – developed during the exposure to concrete and physical disgust stimuli (also termed core or physical disgust in the corresponding literature; see^7,8,14^) – could be extended into the sociomoral disgust domain (e.g., unfairness^3,72^), we applied the VIDS pattern to another independent fMRI dataset during which participants were confronted with a series of unfair offers in an Ultimatum Game task (study 8, *n*=43; Supplementary Methods and Supplementary Table 1). Specifically, we calculated forced-choice classification accuracies between high (average of unfairness level 4 and 5), moderate (unfairness level 3) and low (average of unfairness level 1 and 2) unfairness based on the pattern responses. Noteworthy, disgust is not the only emotion evoked in response to unfairness, other negative emotions might also be involved^3^, we thus compared the performance of PINES and VIFS with VIDS in the sociomoral context. Next, to further examine how specific the VIDS is for unfairness as compared to other social contexts (e.g., pain empathy), we tested the performance of VIDS on another independent dataset (study 9, *n*=238, fMRI brain responses to physical pain and respective non-painful control images as well as to painful faces and respective non-painful control images^49^; Supplementary Table 1). Pattern response was estimated as explained above.

### Reporting summary

Further information on research design is available in the Nature Portfolio Reporting Summary linked to this article.

## Data availability

fMRI data are available at https://figshare.com/articles/dataset/Discovery_dataset_disgust/22827974 and https://figshare.com/articles/dataset/validation_dataset_disgust/22841117.

## Code availability

Data were analyzed using CANLab neuroimaging analysis tools available at https://github.com/canlab and https://github.com/ganxianyang/fMRI-studies/tree/main/Subjective_disgust_experience_signature.

## Acknowledgments

We thank David Coynel and Dominique J.-F. de Quervain as well as Shengdong Chen for sharing their data. We also thank the CANLab for providing the PINES signature and the PINES holdout dataset. In addition, we thank Xiaohong Tian and Qinxia Xie (both majored in pharmacy) who provided us with the essential background knowledge on how to calculate the concentration of different gustatory liquids and the appropriate medical-grade equipment for the gustatory experiment. Any opinions, findings, conclusions or recommendations expressed in this publication do not reflect the views of the Government of the Hong Kong Special Administrative Region or the Innovation and Technology Commission. This work was supported by the National Natural Science Foundation of China (Grants No. 32250610208, 82271583, 32300862), National Key Research and Development Program of China (Grant No. 2018YFA0701400), the China MOST2030 Brain Project (Grant No. 2022ZD0208500), and the Fundamental Research Funds for the Central Universities (SWU2309733).

## Author contributions

X.Y.G., F.Z., and B.B. conceived and designed the experiment. X.Y.G., F.Z., and B.B. analyzed the data and were responsible for interpretation of data. T.X., X.B.L., R.Z., Z.H.Z., X.Y., X.Q.Z., F.W.Y., J.L.L. and R.F.C. provided important suggestions during formal analysis. X.Y.G., R.Z., T.X., and L.W. conducted the experiment. X.Y.G. and R.F.C. were responsible for visualization. X.Y.G. and B.B. drafted the manuscript, F.Z., J.J.Y., and D.Z.Y. provided feedback and revised the manuscript. B.B. supervised the project and acquired the funding. All authors meet the four ICMJE authorship criteria and were responsible for revising the manuscript, approving the final version for publication and for accuracy and integrity of the work.

## Additional information

Extended data is available for this paper at Supplementary information The online version contains supplementary material available at Correspondence and requests for materials should be addressed to Benjamin Becker.

---

**Extended Data Fig. 1.**
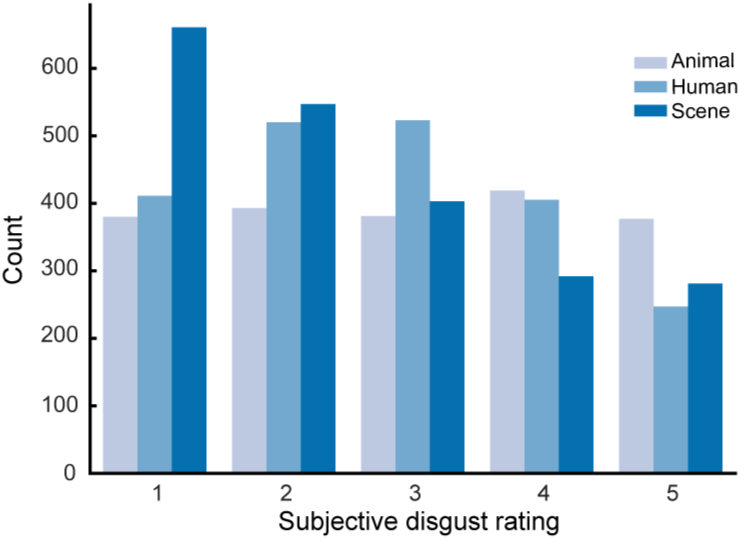
The distribution of subjective disgust ratings for each category (i.e., animal, human, and scene), respectively.

**Extended Data Fig. 2.**
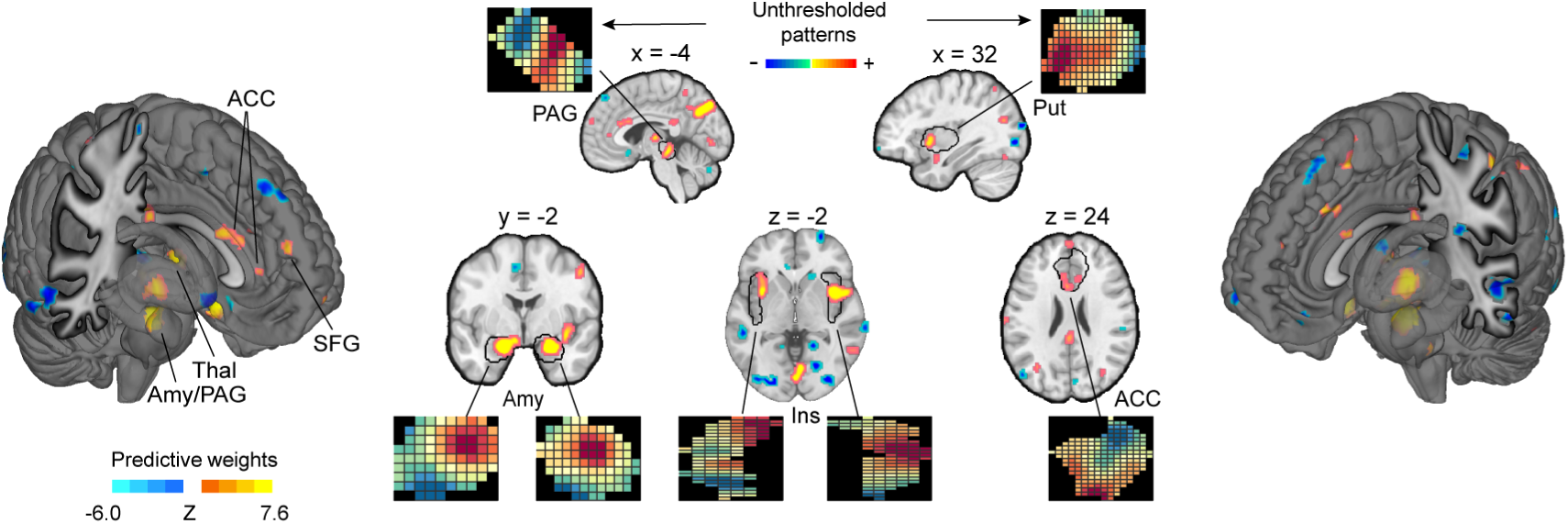
The spatial topography of the unthresholded patterns in some anatomical regions of interest (ROIs). This panel illustrates the VIDS pattern thresholded using a 10,000-sample bootstrap procedure at *q*<0.05, FDR corrected. Inserts show the spatial topography of the unthresholded patterns in some anatomical ROIs. ACC=anterior cingulate cortex, Amy=amygdala, Ins=insula, PAG=periaqueductal gray, Put=putamen, SFG=superior frontal gyrus, Thal=thalamus.

**Extended Data Fig. 3.**
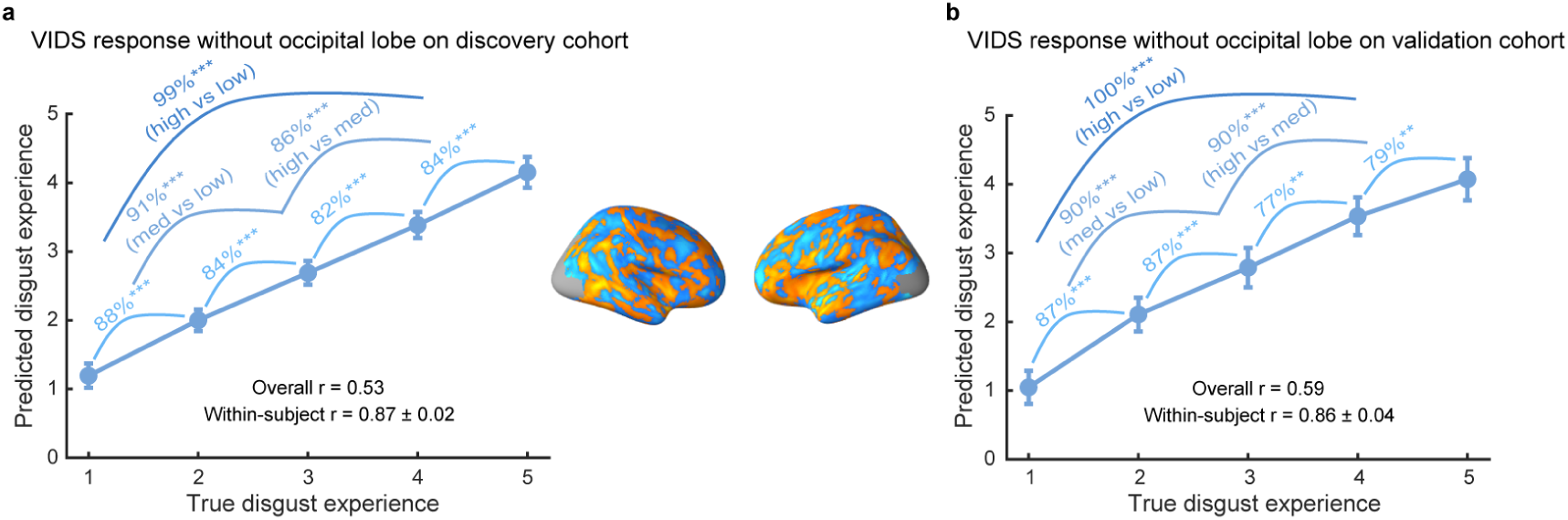
VIDS pattern response without occipital lobe. **a**, The predicted disgust ratings compared to the true ratings for the cross-validated discovery cohort (*n*=78). **b**, The predicted disgust ratings compared to the true ratings for the independent validation cohort (*n*=30). Accuracies reflect forced-choice comparisons. Two-sided binomial tests were used to test whether the classification accuracies were higher than the chance-level. *r* indicates the Pearson correlation coefficient between predicted and true ratings. Error bars reflect the standard error of the mean. ** *P*<0.01 and *** *P*<0.001.

**Extended Data Fig. 4.**
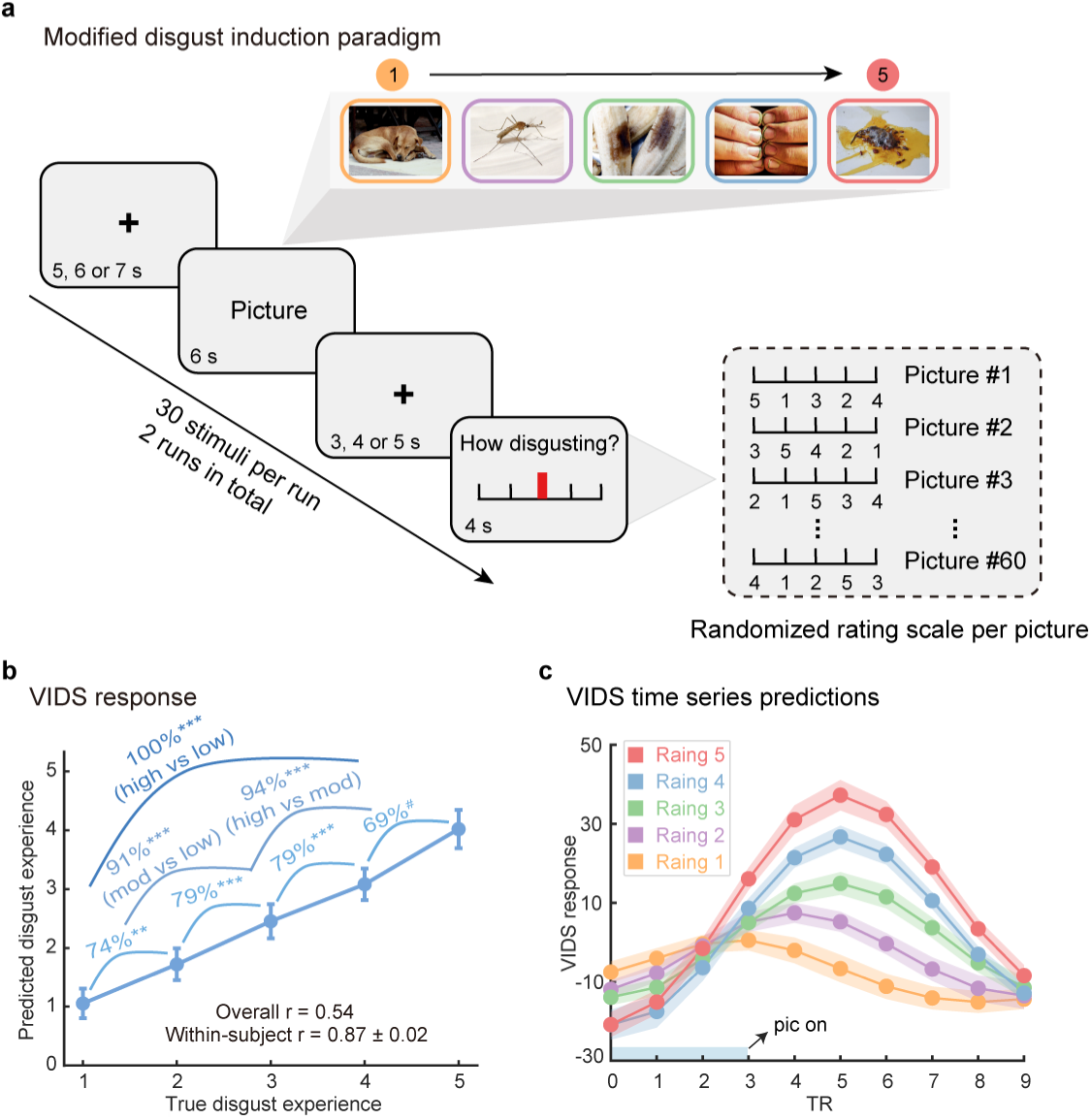
The VIDS tracks disgust experience independent of the motor responses. **a**, The modified disgust induction paradigm included a jittered period between the stimulus and the rating, and the rating numbers were provided in a randomized order. This allowed to better uncouple the motor and emotional response. Example stimuli from the DIRTI disgust images database^59^ (note that all pictures from the DIRTI are subject to creative commons license CC BY-NC 4.0). **b**, Predicted disgust experience (subjective ratings; mean±SE) compared to actual disgust ratings using data from the modified disgust induction task (acquired in *n*=34 individuals). Accuracy provided for forced-choice comparisons. *P* values based on two-sided independent binomial tests. *r* indicates Pearson correlation coefficient between predicted and true ratings. **c**, Averaged peristimulus plot (mean±SE) of the VIDS response using data from the modified disgust induction task at every repetition time (TR; 2s) for each disgust intensity rating separately. ^#^=0.0501, ** *P*<0.01, *** *P*<0.001. Error bars and shaded regions indicate SEs.

**Extended Data Fig. 5.**
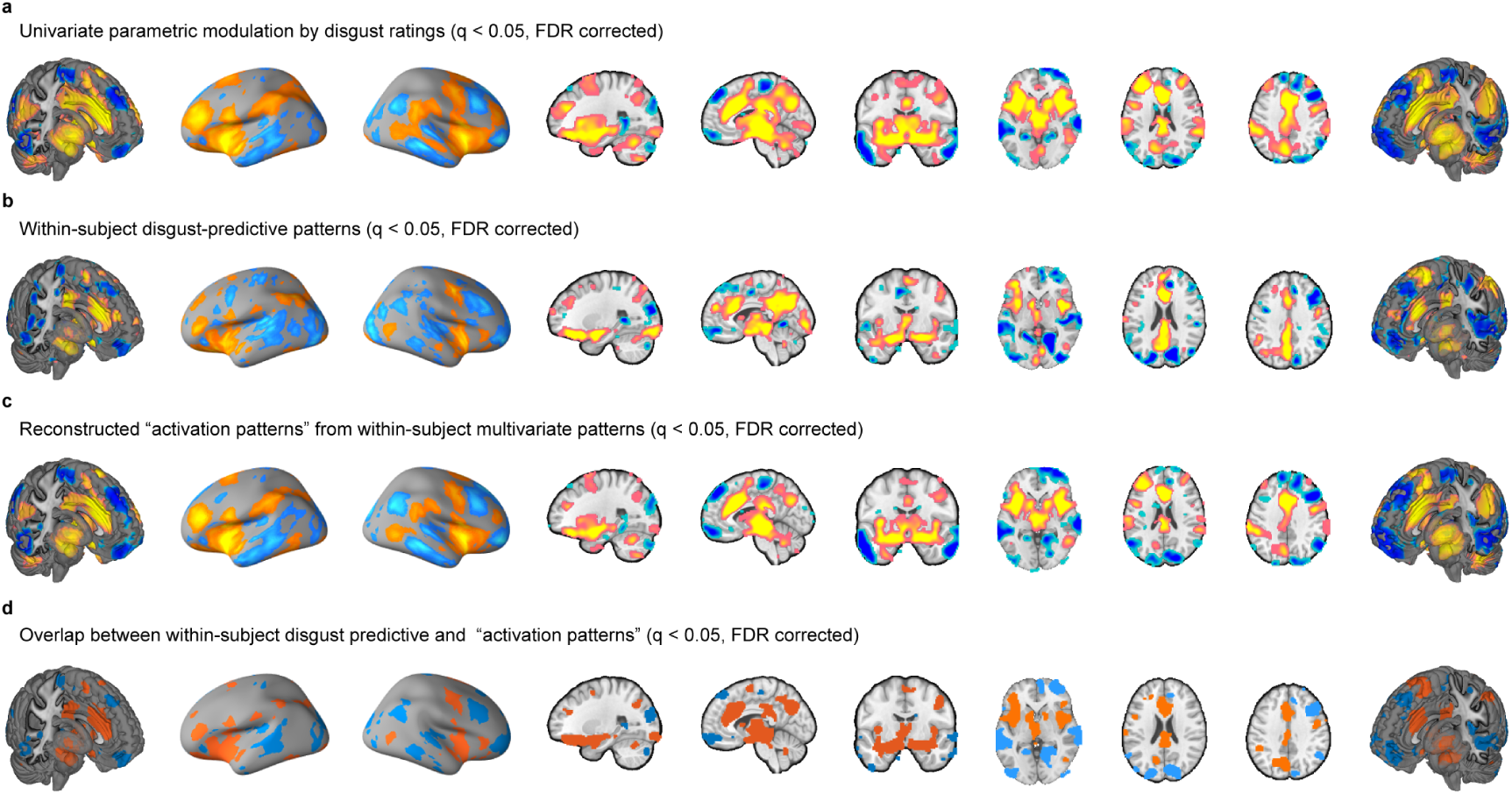
Subjective experience of disgust is associated with and predicted by distributed brain regions. **a**, The univariate parametric effects of disgust ratings. **b**, Multivariate patterns trained on individual subjects and depicts brain regions consistently predictive of subjective disgust across participants. **c**, Thresholded transformed ‘activation patterns’ from within-subject disgust-predictive patterns. **d**, Overlapping (i.e., from a conjunction analysis) brain regions between (**b** and **c**). Hot color indicates positive associations (**a** and **c**) or weights (**b**) whereas cold color indicates negative associations (**a** and **c**) or weights (**b**).

**Extended Data Fig. 6.**
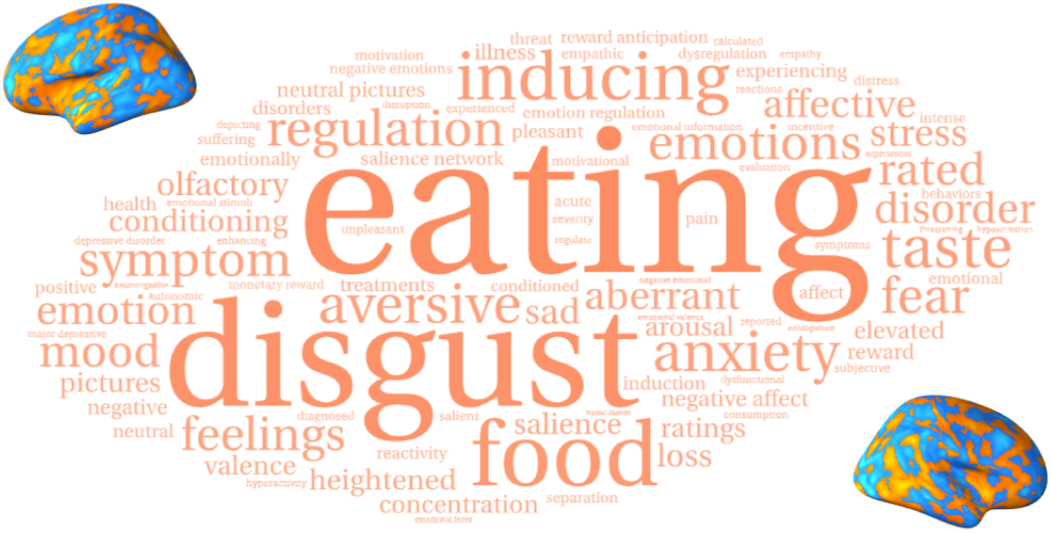
Neurosynth functional decoding of the unthresholded VIDS. Here, the 100 most strongly correlated terms were displayed, with a larger font size indicating a larger Pearson correlation coefficient.

**Extended Data Fig. 7.**
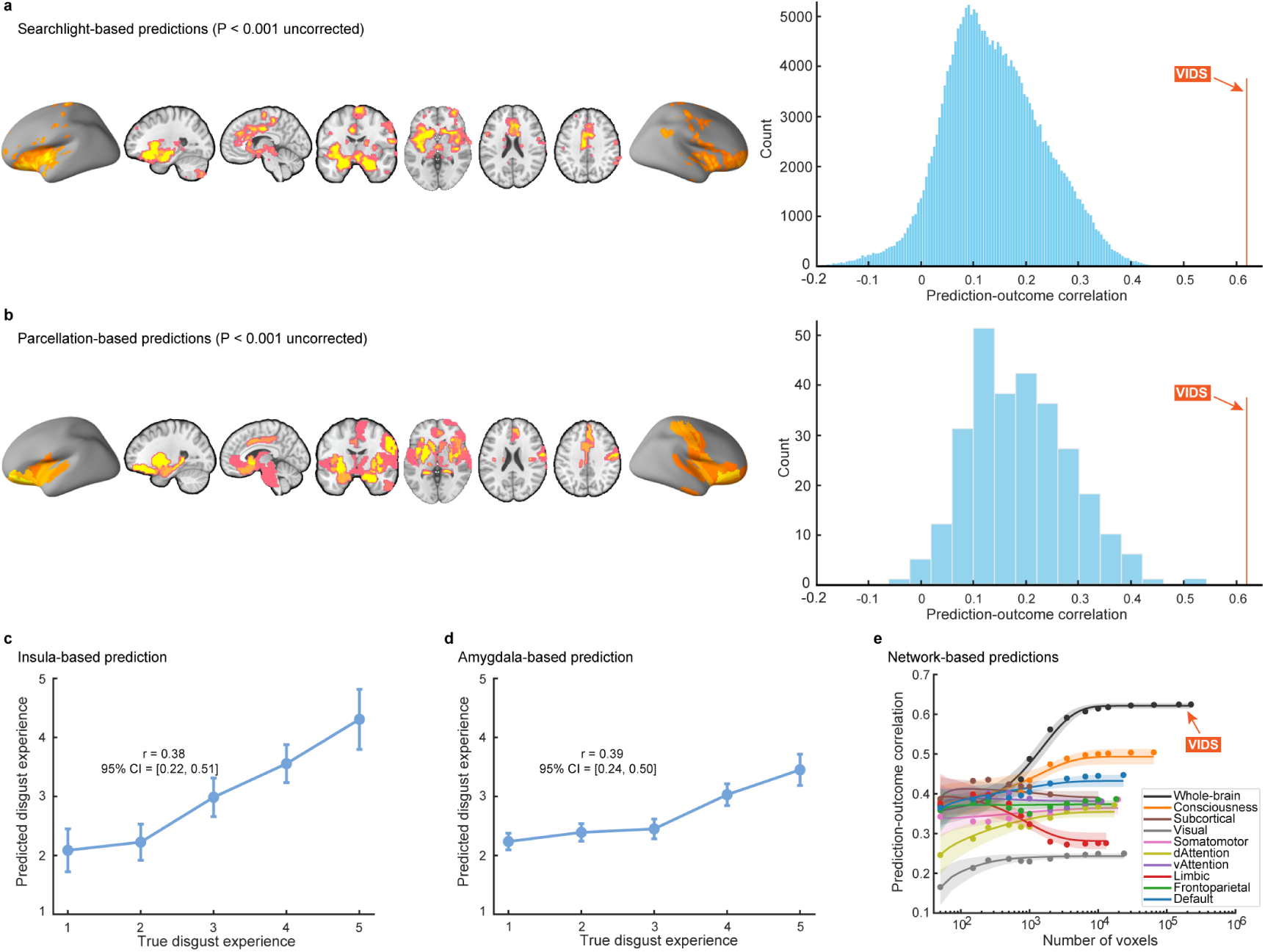
Predictions of models trained on discovery cohort on validation cohort. **a,b**, Brain regions that can significantly predict subjective disgust revealed by searchlight- and parcellation-based analyses, respectively. Histograms: Predictions (correlations) from searchlights and parcellations, respectively. The orange line indicates the prediction-outcome correlation from VIDS. **c,d**, Predictions (mean±SE) from insula- and amygdala-based prediction analyses, respectively. Error bar indicates standard error of the mean; *r* indicates overall (between- and within-subjects; i.e., *n*=149 pairs) prediction-outcome Pearson correlation coefficient. **e**, The information about subjective experience of disgust is distributed across multiple systems. Model performance was evaluated as increasing numbers of voxels/features (x-axis) were used to predict subjective disgust in different regions of interest including the entire brain (black), consciousness network, subcortical regions or large-scale cerebral networks. The y-axis denotes the prediction-outcome correlation. Colored dots indicate the mean correlation coefficients, solid lines indicate the mean parametric fit and shaded regions indicate standard deviation.

**Extended Data Fig. 8.**
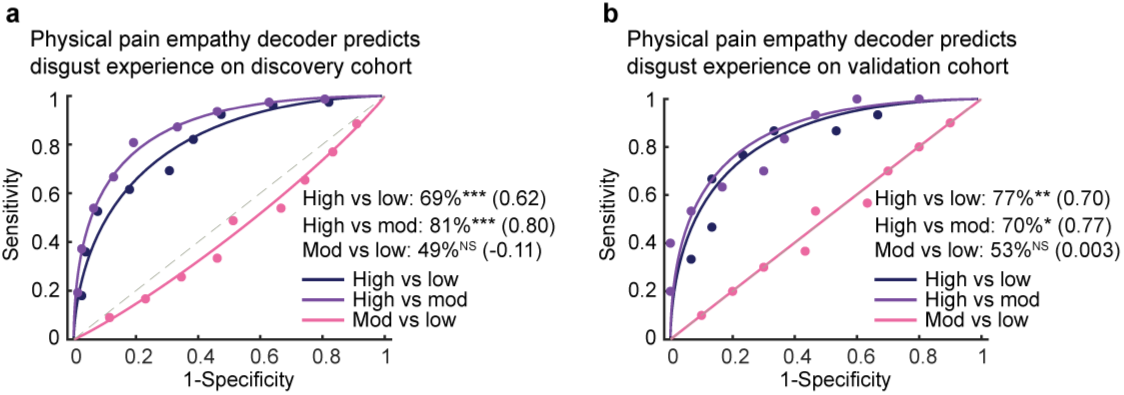
Physical pain empathy decoder predicts disgust experience. **a**, The physical pain empathy decoder could predict high versus low and high versus moderate disgust, nonetheless, it fails to discriminate moderate versus low disgust in the discovery cohort. **b**, The classification results of the physical pain empathy decoder in the validation cohort, which replicates the findings as shown in (**a**). * *P*<0.05, ** *P*<0.01, *** *P*<0.001, NS not significant.

**Extended Data Fig. 9.**
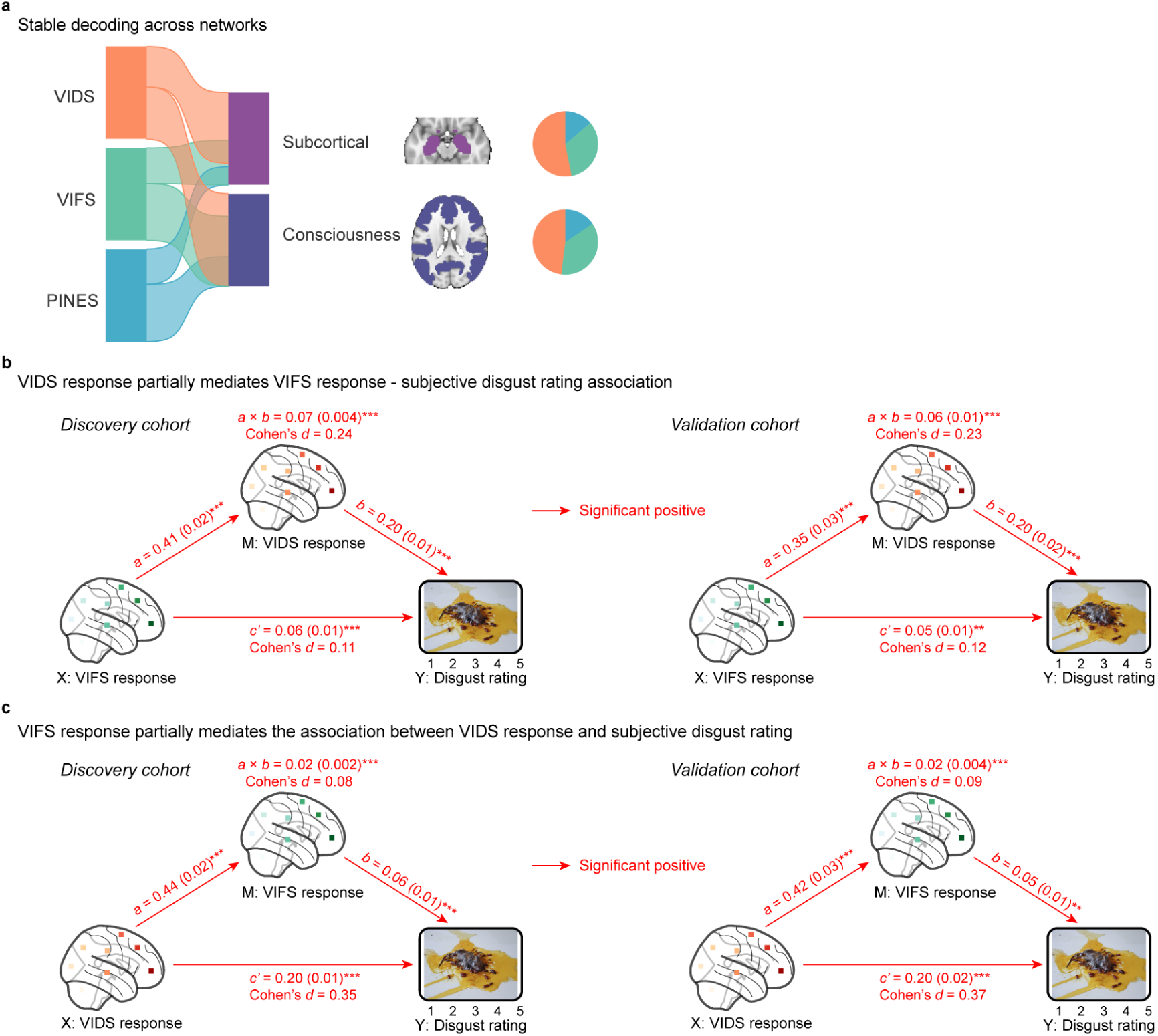
Comparing VIDS, PINES, and VIFS. **a**, River plots displaying spatial similarity (calculated as cosine similarity) between the stable decoding maps and the subcortical as well as consciousness network. Ribbons are normalized by the max cosine similarity across networks. Stable decoding models were thresholded at FDR *q* < 0.05 and positive voxels were retained only for similarity calculation and interpretation. Ribbon locations in relation to the boxes are arbitrary. Pie charts show relative contributions of each model to each network (i.e., percentage of voxels with highest cosine similarity for each map). **b**, The multilevel mediation analytic results showing that VIDS response partially mediates the association between VIFS response and subjective disgust rating in both discovery and validation cohorts. **c**, The VIFS response plays a partial mediation role in the effect of VIDS response on the disgust rating. **b,c**, The example picture from the DIRTI disgust images database^59^ (note that all pictures from the DIRTI are subject to creative commons license CC BY-NC 4.0); and the mediation analysis examines whether the observed covariance between the independent variable (X) and the dependent variable (Y) can be explained by the third variable (M, also mediator), details see Methods section. ** *P*<0.01, *** *P*<0.001 (bootstrap tests with 10,000 samples; two-sided).

**Extended Data Fig. 10.**
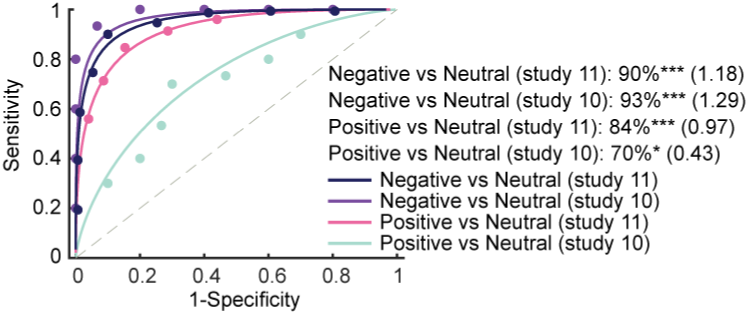
VIDS predicts negative/positive versus neutral emotion. The VIDS reacted somehow to negative/positive versus neutral emotion. * *P*<0.05, *** *P*<0.001.

