## Supplementary information for "Rotten to the core – a neurofunctional signature of subjective core disgust generalizes to oral distaste and socio-moral contexts"

**Supplementary Methods**

**Participants in the modified disgust induction experiment (Study 3)**

The modified disgust induction task included thirty-five healthy participants recruited from the UESTC. Data from one female participant were excluded due to excessive head movement (>3mm) during fMRI scanning, leading to a final sample of *n*=34 participants (15 females; mean±SD age=23.50±2.27 years). Enrollment criteria were identical to study 1. Informed consent was obtained before the experiment, the experimental protocol was approved by the UESTC Ethics Board and in line with the latest revision of the Declaration of Helsinki. Participants were reimbursed 60 RMB.

**Stimuli and paradigm used in the modified disgust induction experiment**

The modified disgust induction task included 60 pictures, all of which were from the Disgust-related Images (DIRTI) database^1^, distributed over 2 runs (with 30 stimuli per run). Forty out of 60 stimuli overlapped with the stimuli used in the discovery and validation cohorts, 20 new stimuli were added to explore whether the signature generalizes robustly also across stimuli. Stimuli were presented using E-Prime 3.0. Participants were instructed to pay attention to the pictures and naturally experience the induced emotion. Each trial consisted of a 6s presentation of the picture followed by a jittered fixation-cross (3, 4, or 5s) separating the stimuli from the rating period. Participants then were requested to report within 4s the level of disgust they experienced for the stimuli using a 5-point Likert scale with 1 indicating neutral/slightest disgust and 5 indicating most strongly disgust. Note that the order of rating 1 to rating 5 was no longer ascending but in a randomized order presented for each rating to allow a further separation of the emotional and motor response. Finally, there was a jittered fixation-cross epoch (5, 6, or 7s) before the presentation of the next picture (Extended Data Fig. 4a). All participants reported ‘1-4’ in their responses while 2 out of 34 participants did not report the rating ‘5’.

**MRI data acquisition for the modified disgust induction experiment**

MRI data were acquired on a 3T system (GE MR750, General Electric Medical System, Milwaukee, WI, USA). Structural images were collected using a high-resolution T1 spoiled gradient recall (SPGR) sequence (repetition time=8ms, echo time=3ms, flip angle=8°, field of view=256×256mm, voxel size=1×1×1mm, acquisition matrix=256×256, 176 slices, slice thickness=1mm) and were used for improving spatial normalization and excluding participants with apparent brain pathologies. Functional images were acquired with a T2*-weighted echo planar imaging (EPI) sequence (repetition time=2000ms, echo time=30ms, flip angle=90°, field of view=200×200mm, voxel size=3.125×3.125×3.8mm, resolution=64×64, 36 slices, slice thickness=3.8mm, no gap).

**fMRI data preprocessing and first-level fMRI analysis for the modified disgust induction experiment**

Preprocessing and subject-level GLM analysis for the modified disgust induction task were identical to the discovery cohort.

**MRI data acquisition in the generalization cohort (Study 4)**

Data were acquired on a Siemens Trio 3T MRI system (Magnetom Trio, Siemens, Erlangen, Germany) with a gradient echo planar imaging sequence (32 axial slices, TR/TE=2s/30ms, FA=90°, matrix size=64×64, FOV=220×220mm, voxel size=3.4×3.4×3mm, and 386 functional volumes). High-resolution structural images were acquired for registration purposes using a T1-weighted magnetization-prepared rapid gradient-echo (MP-RAGE) sequence (TR/TE=1900ms/2.52ms, FA=9°, FOV=256×256mm, slices=176, thickness=1.0mm, and voxel size=1×1×1mm).

**Determining brain regions associated with and predictive of subjective disgust on the individual level**

We examined regions involved in the subjective experience of disgust on the individual level by employing convergent univariate and multivariate approaches. We initially determined regions that were predominately associated with subjective disgust ratings by means of a one-sample *t* test on the first-level univariate parametric modulation beta maps. Next, to localize brain regions that were predictive of and associated with disgust ratings separately as well as brain regions showing shared contributions, multivariate analyses were employed. Briefly, we evaluated the consistency of each weight for every voxel in the brain across within-subject multivariate classifiers (developed on single-trial data) using a one-sample *t* test. The thresholded map (*q*<0.05, FDR corrected for multiple comparisons) revealed consistent disgust-predictive brain regions across subjects. Specifically, we performed a prediction analysis (linear SVR with *C*=1) for each subject in the discovery cohort separately using their single-trial data (10×10-fold cross-validated) and only included participants whose disgust ratings could be significantly predicted by their brain data (evaluated by prediction−outcome Pearson correlation; *n*=72, from 78 participants). Of note, similar results were found when including the entire sample in the analysis. Moreover, we transformed the within-subject patterns into ‘activation patterns’ to identify significant brain regions exhibiting consistent disgust-related effect (thresholded at *q*<0.05, FDR corrected).

**Examination of the physical pain empathy decoder for predicting subjective disgust experience**

Zhou et al. developed a whole-brain signature for (physical) vicarious pain that could accurately discriminate brain responses towards visual stimuli depicting physical pain infliction versus corresponding matched non-painful control stimuli^2^. Given that both, pain empathy and disgust are traditionally considered as prototypical affective responses linked to the insular cortex, we utilized this signature to further determine the specificity of the disgust decoder for functions closely related to the insula. To this end, we employed additional predictions using the (physical) vicarious pain pattern to the disgust discovery and validation datasets. We specifically tested whether the pattern could accurately predict different levels of disgust experience. Estimation of the pattern response was computed using the dot-product of each vectorized brain activation image with the targeted pattern, thus yielding a continuous, scalar value.

**Participants in the gustatory experiment (study 7)**

To test whether the visual disgust decoder (i.e., VIDS) generalizes to the gustatory modality, we designed an fMRI paradigm using gustatory disgust and neutral stimuli (orally applied liquids). The paradigm was utilized to acquire data in an independent dataset (study 7). The gustatory experiment included thirty healthy participants (15 females; mean±SD age=24.27±2.26 years) recruited from the UESTC. Enrollment criteria were identical to study 1. Informed consent was obtained before the experiment, the experimental protocol was approved by the UESTC Ethics Board and was in line with the latest revision of the Declaration of Helsinki. Participants were reimbursed 90 RMB.

**Chemosensory stimuli and paradigm used in the gustatory experiment**

Consistent with previous behavioral and neuroimaging studies^3,4^, disgusting liquids were chosen from different concentrations of salty sodium chloride solutions (NaCl; 0.1, 0.5, 1 mol) and sour citric acid solutions (C_6_H_8_O_7_; 0.002, 0.01, 0.05, 0.1 mol). We used bitter gourd beverages (96% of the content is bitter gourd ingredient; purchased from e-commerce stories) as the bitter condition (for similar approaches, see ref.^5^). In keeping with earlier studies^4,6,7^, artificial saliva (2.5mM NaHCO_3_ and 25mM KCl; diluted with 30%, 50%, and 80% water) rather than water was selected as the neutral taste. Adopting evaluated disgust thresholding procedures^3,4^, we utilized individualized liquids as disgusting stimuli for each subject. Specifically, prior to the fMRI experiment, participants were required to participate in a behavioral taste experiment during which a variety of liquids was administered (bitter gourd beverages, three concentrations of salty solutions, four concentrations of sour solutions, and three concentrations of artificial saliva solutions, 1ml per liquid). After consuming each solution participants rated their experience using 5-point Likert scales in terms of subjective disgust experience (how disgusting? ‘not at all’ to ‘very disgusting’) and taste intensity (intensity of the taste? ‘barely detectable’ to ‘strongest imaginable’). For each subject, the disgusting liquid with a rating closest to ‘very disgusting’ and the neutral liquid with a rating close to ‘not at all disgusting’ were chosen. Specifically, for bitter liquid (chosen for 13 participants), mean±SD subjective disgust rating=4.54±0.52; for sour liquid (6 participants), mean±SD subjective disgust rating=4.50±0.55; for salty liquid (11 participants), mean±SD subjective disgust rating=4.82±0.40, suggesting that the chosen liquids elicited a high-level of subjective disgust experience. Moreover, a one-way ANOVA revealed that the main effect of substance type was not significant (*F* (2, 27)=1.27, *P*=0.297), indicating similar disgust levels across the chosen liquids. The neutral liquid was rated with mean±SD subjective disgust rating=1.07±0.25 (in all 30 participants) indicating a strong difference between the conditions in terms of subjective disgust experience. Finally, the intensities of the chosen disgusting stimuli were high (mean±SD=4.37±0.61) and congruent with previous studies^3,4,8^.

In the fMRI experiment, we employed similar gustatory procedures as described in earlier studies^6,7^. During fMRI the individuals were lying supine in the MRI system and received visual instructions, while an experimenter standing beside the MRI system in the MRI room received auditory instructions (Fig. 1g) to facilitate a precise timing of the administration. Upon the time-logged instructions the liquid solutions were administered by the experimenter using 29.5 cm-long medical-grade disposable plastic pipettes. The timing of each trial, auditory instructions, and visual instructions were presented and synchronized with the MRI system using E-Prime 3.0. At the beginning of each trial, participants were presented a text on the screen instructing them to open their mouth, the experimenter simultaneously received an auditory instruction to deliver 1ml of disgusting/neutral liquid onto the participant's tongue using the pipette and continuously over a 5s period. Next, a ‘taste’ instruction was presented on the screen for 8s, and during this period participants were required to hold the liquid in their mouth and taste it, until a ‘swallow’ instruction (5s) was shown on the screen indicating to swallow the liquid. To minimize carry-over effects between subsequent trials, the experimenter then received another auditory instruction to deliver 2ml of water for rinsing in the participant’s mouth via a separate pipette (5s). Next a ‘rinse’ instruction was shown to the participant to indicate them to rinse their mouth (6s), followed by a ‘swallow’ instruction (5s). Trials were separated by a jittered fixation-cross epoch (4, 5, or 6s). The current gustatory experiment included two disgust runs and two neutral runs, with 6 trials per run. The order of the four runs was either ‘disgust-disgust-neutral-neutral’ or ‘neutral-neutral-disgust-disgust’, and counterbalanced across participants. Before scanning, the participants were familiarized with the entire procedure using water and were trained to undergo the procedure with minimal head motion.

**MRI data acquisition for the gustatory experiment**

Imaging data acquisition for the gustatory experiment was identical to the modified disgust induction task (study 3).

**fMRI data preprocessing for the gustatory experiment**

Preprocessing for the gustatory experiment was identical to the discovery cohort. Despite strong gustatory sensations as well as swallowing of gustatory stimuli, the head movements for 116 out of 120 functional runs were within 1mm, with the remaining four runs within 1.5mm.

**First-level fMRI analysis for the gustatory experiment**

GLM analyses were implemented in SPM following standardized preprocessing of the data. The four runs of the fMRI task were concatenated for each participant. We modeled liquid delivery, tasting conditions for disgust and neutral liquids, swallowing of these tasting conditions, rinsing of these tasting conditions, and swallowing of the rinsing water in the GLM model with the onsets time-logged to corresponding conditions. Task regressors were convolved with the canonical hemodynamic response function and a high-pass filter of 128s was applied. Nuisance variables encompassed: (a) ‘dummy’ regressors representing each run (intercept for each run); (b) the six estimated head movement parameters (X, Y, Z, roll, yaw, and pitch), their squares, their derivatives and squared derivatives for each run (24 columns in total); (c) vectors indicating motion outlier time points. For each subject, we included the beta images related to the taste (as well as swallow) of disgust and neutral liquids in the following prediction analyses.

**Ultimatum Game paradigm in study 8**

The study 8 dataset included in the present paper is part of a larger study during which participants completed a modified version of the Ultimatum Game (UG) paradigm^9^ with concomitant fMRI. During the UG participants were in the role of the responder who received a series of offers from the proposer. In line with the standard version of the UG, the paradigm included 20 trials with offers of distributions of a fixed sum (100RMB per trial). The unfairness of the offer followed a randomized but uniform distribution such that each participant would be presented with the full range of offers ranging from fair (50RMB versus 50RMB) to very unfair (92.5RMB versus 2.5RMB). Participants were informed that if they accepted the monetary offer, the money would be divided between them and the proposer according to the offer, however, if they rejected the offer, neither side would receive any money. The modified version included additional trial-wise ratings regarding: (1) the amount of money (within a range of 2.5RMB to 50RMB, 20 offers) they would expect; (2) the prediction of the emotional experience of the anticipated offer, and (3) report of the actual emotional experience in response to the offer. In the context of the current study, we focused on the classic part of the UG during which the participant was presented with an offer by the proposer and decided whether to accept or reject the offer^3^. Results of the modification (behavioral and neural activity related to the expectation and outcome) were part of an independent study (not presented here). Each task trial began with a fixation cross presented for a jittered interval of 2s followed by the presentation of expected offers and ratings, actual offer presentation, and decision periods (mean duration for each task trial=14s). In line with Chapman et al., we focused on the first 20 trials of the monetary distribution offers^3^ and further categorized them into 5 levels of unfairness (i.e., unfairness level 1 to 5, see Supplementary Table 5 for details).

**MRI data acquisition, preprocessing, and first level analysis in study 8**

Imaging data acquisition was identical to the discovery and validation cohorts. All fMRI images were preprocessed and analyzed using standard procedures in SPM12 (Statistical Parametric Mapping; http://www.fil.ion.ucl.ac.uk/spm/; Welcome Trust Centre for Neuroimaging). The first 5 volumes of each functional time series were discarded to allow for T1 equilibration. Remaining images were corrected for acquisition time delay, realigned to correct for head motion, unwarped for magnetic field inhomogeneities correction, and co-registered with the T1-weighted structural image. After that the images were normalized to Montreal Neurological Institute (MNI) standard space (interpolated to 2×2×2mm voxel size) using the segmentation parameters from the anatomical images, and were then spatially smoothed using an isotropic Gaussian kernel with full-width at half-maximum (FWHM) of 8mm. The 20 pairs of offers were divided into 5 levels of unfairness (see Supplementary Table 5), and included as separate regressors in the GLM model with the onsets time-logged to the actual presentation of the offer. The six head motion parameters were included as covariates of no interest.

**Datasets to validate a potential contribution of arousal**

To explore whether VIDS may capture general and non-specific arousal which is inherent to several strong emotional experiences (including e.g., fear and disgust), we tested the performance of the VIDS on two additional independent datasets. Both studies presented neutral (low arousing) and strongly negative and positive (high arousing) pictures to healthy individuals during fMRI (study 10, *n*=30, fMRI brain responses to negative, positive, and neutral pictures; and study 11, *n*=150, fMRI brain responses to negative, positive and neutral pictures; Supplementary Table 1). As for the estimation of the pattern response, we computed the dot-product of each vectorized brain activation image with the VIDS pattern, thus yielding a continuous, scalar value.

**Supplementary Results**

**VIDS requires, but not fully depends on, the visual cortex**

To examine to which extent the performance of VIDS may depend on the contribution of the visual cortex – which might encode emotional experience or aspects of high-level visual processing related to emotion^10,11^ - the disgust decoder was re-trained excluding the occipital lobe. Results showed that the overall prediction-outcome correlations (the discovery cohort, cross-validation *r*=0.53, 95% CI=[0.45, 0.59]; the validation cohort, *r*=0.59, 95% CI=[0.46, 0.67]), within-subject correlation between predicted and true disgust ratings (the discovery cohort, *r*=0.87±0.02; the validation cohort, *r*=0.86±0.04) and classification accuracies were largely maintained (see Extended Data Fig. 3a,b for details). This indicates that disgust-predictive neural representations are partly embedded in the visual cortex but the contribution of visual cortical patterns is comparably small.

**VIDS indeed decodes the experience of disgust rather than the mapping of disgust onto responses**

To exclude that the VIDS might partly decode the mapping of disgust onto motor responses (rather than the experience of disgust) – because the interval between the picture presentation stage and button pressing stage was fixed in the original disgust induction paradigm (in discovery and validation cohorts, Fig. 1b) - we conducted an additional fMRI experiment that uncouples the motor and emotional response (study 3). The modified disgust induction paradigm included an additional jitter between picture presentation and motor responding stage; and the order of rating 1 to rating 5 was no longer ascending but in a randomized fashion presented per trial (details see Extended Data Fig. 4a). Applying the predictive model - with no further model fitting – to the dataset from the modified disgust induction task revealed comparable high prediction-outcome correlations (within-subject *r*=0.87±0.02, average RMSE=1.56±0.13; overall prediction-outcome *r*=0.54, 95% CI=[0.42, 0.63]; Extended Data Fig. 4b), indicating a sensitive and robust neurofunctional signature for subjective disgust experience. Furthermore, the VIDS response accurately classified high (average of rating 4 and 5) versus moderate (rating 3) and moderate versus low (average of rating 1 and 2) disgust with 91%-94% accuracy (Cohen’s *d*=1.31–1.38), and high versus low with 100% accuracy (Cohen’s *d*=2.95). Moreover, the VIDS response could distinguish each successive pair of disgust rating levels (e.g., rating 1 versus rating 2) with high accuracy (significantly higher than 50% chance-level; except rating 5 versus 4, with accuracy being 69% and *P*=0.0501) (Extended Data Fig. 4b).

To further determine the specificity of the VIDS to the experience of disgust (rather than anticipation or cognitive evaluation), we explored the VIDS reactivity over the stimulus presentation interval covering the pre-stimulus interval and onset of the stimulus in the dataset from the modified disgust induction task. In line with the standard hemodynamic model, the VIDS response started approximately 4s following the picture onset and increased during 6-12s (Extended Data Fig. 4c), suggesting that the VIDS captures disgust experience rather than expectation (pre-stimulus) or cognitive evaluation (response-reporting) of disgust.

Together with similar findings observed in the discovery and validation cohorts, these results suggest a robust and generalizable neural signature that captures subjective disgust experience across different paradigms, stimuli, populations, and scanning protocols.

**Brain regions associated with and predictive of subjective disgust on the individual level**

To identify regions involved in the subjective experience of disgust on the individual level, convergent univariate and multivariate approaches were employed. First, we performed a one-sample *t* test on the first-level parametric modulation beta maps to determine brain regions that increased activity in association with the subjective experience of disgust (i.e., the higher the ratings the higher functional activation in the respective brain regions). Results revealed that subjective disgust was correlated with activation in a broad network spanning multiple cortical and subcortical systems, with increasing activation in a bilateral network encompassing the insula, amygdala, cingulate cortex (covering anterior, middle and posterior parts), thalamus, basal ganglia (e.g., putamen, caudate and globus pallidus), midbrain regions including the PAG, supplementary motor area (SMA) and occipital-temporal-parietal regions. In contrast, negative associations with disgust ratings were observed in the middle frontal gyrus, superior frontal gyrus, and middle temporal gyrus (thresholded at FDR *q*<0.05 corrected for multiple comparisons; Extended Data Fig. 5a).

Next, we validated and extended the univariate results by multivariate models in several ways. We first performed a one-sample *t* test (treating participant as a random effect) on the weights obtained from within-subject multivariate predictive models (for details, see ‘Supplementary Methods’). Similar to the univariate maps, within-subject predictive models (backward models) included consistent weights in brain regions spanning multiple large-scale cortical and subcortical systems that largely overlapped with the disgust regions as determined by the univariate parametric modulation approach (Extended Data Fig. 5b; *q*<0.05, FDR corrected).

To provide a more direct comparison between univariate and multivariate results, we further calculated within-subject reconstructed ‘activation patterns’ (or ‘structure coefficients’), which map each voxel to the overall multivariate model prediction^12^. Extended Data Fig. 5c shows the thresholded transformed ‘activation patterns’ from the within-subject disgust-predictive patterns. As shown in Extended Data Fig. 5c, the thresholded ‘model activation patterns’ display an overlap with the univariate and multivariate patterns (Extended Data Fig. 5a,b), confirming that the subjective experience of disgust is encoded in a distributed neural representation encompassing cortical and subcortical regions. The overlap of the multivariate predictive models (i.e., conjunction between within-subject model weights and model encoding maps; Extended Data Fig. 5d) revealed that most significant model weights in the multivariate models also encoded model information (i.e., showed significant ‘model activation patterns’). The broad conclusion is that the neural representation of human disgust is not limited to a single or a set of focal regions (e.g., the insula), but rather encoded in a broad set of regions spanning cortical and subcortical systems.

**Meta-analytic decoding of the unthresolded VIDS pattern based on Neurosynth also supported VIDS as a biologically plausible model of disgust**

To complement the BrainMap functional characterizations, we further capitalized on the Neurosynth database^13,14^ to functionally decode the unthresholded VIDS pattern (for similar approaches, see ref.^15^). The top 100 terms were displayed with a larger font size indicating a higher convergence (i.e., Pearson correlation coefficient; Extended Data Fig. 6 and Supplementary Table 2). The results revealed terms mainly related to threat-, avoidance-, and salience-related processes (e.g., ‘disgust’, ‘aversive’, ‘salience’); moreover, it also corroborated that the disgust signature captures subjective disgust experience as indicated by terms such as ‘inducing’, ‘feelings’, and ‘experiencing’. Interestingly, terms associated with gustation (e.g., ‘eating’, ‘food’, ‘taste’) were also highly related, which may reflect the oral origin of disgust (for more evidence, see section ’The visual subjective disgust signature predicts gustatory stimulus induced (taste) disgust’). Together, two data-driven approaches (i.e., meta-analytic decoding based on BrainMap and Neurosynth) converge to support VIDS as a biologically plausible model of disgust.

**VIDS is separable from the neural expressions of other functional domains classically associated with overlapping neural systems, i.e., pain empathy**

To further determine the specificity of the VIDS, we tested whether a neurofunctional decoder of pain empathy (often conceptualized as insula-dependent^2,4,16^) can predict disgust (see ‘Supplementary Methods’ for details) in disgust discovery and validation cohorts. As shown in Extended Data Fig. 8a, in the discovery cohort the physical pain empathy decoder could predict high versus low and high versus moderate disgust experience (accuracy: 69%-81%, Cohen’s *d:* 0.62-0.80). However, in contrast to the disgust decoder, it could not differentiate between moderate and low disgust experience (accuracy=49%, Cohen’s *d*=-0.11). Furthermore, these results were replicated in the validation cohort (Extended Data Fig. 8b). Additionally, the overall prediction-outcome and within-subject correlations (discovery cohort: overall *r*=0.17, within-subject *r*=0.32±0.05; validation cohort: overall *r*=0.21, within-subject *r*=0.43±0.09) were much lower as compared to those obtained by the VIDS in both discovery and validation cohorts (Fig. 2b,c). These results reflect that the (physical) pain decoder is not very accurate in differentiating subjective disgust experience, indicating that even if two functions (i.e., disgust and pain empathy) have been strongly related to the insula, on the whole brain level they can still be separated.

**VIDS response mediates the association between subjective disgust and neural reactivity of a non-specific negative affect signature (PINES)**

We performed multilevel mediation analyses to examine the relationship between VIDS response, disgust ratings and PINES response. We first tested whether the VIDS response (*M*) could mediate the relationship between the PINES response (*X*) and subjective disgust rating (*Y*). The results showed that in the discovery cohort the PINES response was positively correlated with the VIDS response (path *a*; β=0.42, SE=0.03, 95% CI=[0.36, 0.48]; *Z*=3.77, *P*<0.001; Cohen’s *d*=0.27), the VIDS response was positively associated with disgust ratings independently of the PINES response (path *b*; β=0.22, SE=0.01, 95% CI=[0.20, 0.24]; *Z*=3.66, *P*<0.001; Cohen’s *d*=0.42) and there was a significant mediation effect (path *a*×*b*; β=0.08, SE=0.01, 95% CI=[0.07, 0.09]; *Z*=3.58, *P*<0.001; Cohen’s *d*=0.18). Furthermore, the total effect was significant (path *c*; β=0.11, SE=0.01, 95% CI=[0.08, 0.14]; *Z*=3.74, *P*<0.001; Cohen’s *d*=0.12), however, the non-mediated effect was not significant (path *c’*; β=0.02, SE=0.01, 95% CI=[-0.004, 0.04]; *Z*=1.61, *P*=0.10; Cohen’s *d*=0.02). The above findings were replicated in the validation cohort, except that there was a significant non-mediation effect (path *c’*; β=0.05, SE=0.02, 95% CI=[0.01, 0.09]; *Z*=2.37, *P*=0.02; Cohen’s *d*=0.07) in the validation cohort. Together, our results indicate that the VIDS response plays a full mediation role in the effect of PINES response on the subjective disgust rating in the discovery cohort, and in validation cohort the VIDS response partially mediated the effect of PINES response on the subjective disgust rating (Fig. 6e).

Additionally, a second multilevel mediation model was constructed with the VIDS response as *X*, the PINES response as mediator (*M*) and subjective disgust rating as *Y*. The results revealed that the PINES could not mediate the association between the VIDS response and disgust ratings in both, discovery and validation cohorts (Fig. 6f), suggesting that the VIDS captures neural representations that are – to a certain extent – more specific to disgust than general negative affect.

**VIDS response mediates subjective disgust induced by fear**

We further employed two multilevel mediation analyses to investigate the relationship between VIDS response, VIFS response and subjective disgust rating. Likewise, we aimed to test whether (1) the VIDS response (*M*) could explain the relationship between the neurofunctional signature of a functionally related emotional response (i.e., subjective fear as represented in the VIFS response, *X*) and disgust rating (*Y*), and whether (2) the VIFS response (*M*) could mediate the association between VIDS response (*X*) and disgust rating (*Y*). The results showed that in the discovery cohort, the VIFS response had a positive association with the VIDS response (path *a*; β=0.41, SE=0.02, 95% CI=[0.37, 0.46]; *Z*=3.70, *P*<0.001; Cohen’s *d*=0.49), the VIDS response was positively correlated with disgust rating (path *b*; β=0.20, SE=0.01, 95% CI=[0.18, 0.22]; *Z*=3.54, *P*<0.001; Cohen’s *d*=0.35) and a significant mediation effect emerged (path *a*×*b*; β=0.07, SE=0.004, 95% CI=[0.06, 0.08]; *Z*=3.47, *P*<0.001; Cohen’s *d*=0.24). Moreover, both path *c* (β=0.14, SE=0.01, 95% CI=[0.12, 0.16]; *Z*=3.78, *P*<0.001; Cohen’s *d*=0.28) and path *c’* (β=0.06, SE=0.01, 95% CI=[0.04, 0.07]; *Z*=3.81, *P*<0.001; Cohen’s *d*=0.11) were significant. Overall, the above results demonstrated that the VIDS response played a partial mediation role in the effect of VIFS response on the disgust rating. Noteworthy, these findings were substantiated in the validation cohort (Extended Data Fig. 9b).

We next constructed an alternative mediation model with VIFS response as *M*, VIDS response as *X* and disgust rating as *Y*. Although a partial mediation effect of the VIFS response was observed, the former model (Extended Data Fig. 9b) was more robust than the latter one (Extended Data Fig. 9c) as indicated by effect sizes that were three times higher (model 1: Cohen’s *d*=0.24; model 2: Cohen’s *d*=0.08) in the discovery cohort. These results were verified in the validation cohort. Taken together, these findings suggested that the VIFS response partially mediated the VIDS response effect on the subjective disgust rating, however, this mediation model was not as robust as the first one.

**VIDS is specific and sensitive to track subjective disgust experience rather than non-specific emotional arousal**

We utilized two independent datasets using positive and negative visual stimuli to determine if the VIDS primarily tracks non-specific emotional arousal. The results (Extended Data Fig. 10) showed that the VIDS could accurately predict negative versus neutral pictures (90%-93% accuracy, and 1.18–1.29 Cohen’s *d*), as well as positive versus neutral pictures (70%-84% accuracy, and 0.43–0.97 Cohen’s *d*). However, the effect sizes were not as large as using VIDS to predict disgust experience. For example, the effect sizes in the high versus low disgust context (i.e., 2.40 in the discovery cohort and 2.92 in the validation cohort, Fig. 6c) were 1.86–6.79 times higher than those observed here. Together, this indicates that the VIDS partly captures high negative/positive emotional arousal while the effect sizes additionally indicate a higher specificity of the VIDS to the specific target emotion (disgust).

The question of to what extent non-specific high emotional arousal contributes to neurofunctional signatures of emotional experiences represents a matter of debate. The effect sizes in our study indicate a considerably higher reactivity to disgust than general negative emotions and this is further validated by comparisons and mediation models encompassing a series of other neurofunctional signatures for high arousal emotional states including vicarious pain, fear, and non-specific negative affect. However, it has also been proposed that (see ref.^17^) using different patterns to discriminate intense emotional experiences from control conditions (e.g., neutral) will yield a high accuracy given the fundamental differences between strong emotional states and neutral conditions which include arousal, salience, autonomous reactivity, etc. However, if we could provide evidence that different brain patterns are highly predictive of their targeted emotional experiences, this would support that these brain representations underlie partly distinguishable affective experiences. Indeed, in the current study, we compared the performance of VIDS, VIFS, and PINES across multi-study datasets (e.g., Fig. 6c,d,e,f, Fig. 7a,b, Fig. 8a,b, and Extended Data Fig. 9b,c; Supplementary Table 4) and together the results suggest that the VIDS represents a neurofunctional representation that is - to a certain extent – distinguishable from the representation associated with the subjective experience of some other affective and motivational states.

**Supplementary Tables**

Supplementary Table 1. Information for each study.

| *Study* | *Dataset ref.* | *Stimuli Types* | *N (female)* | *Mean age (SD)* | *Study location* | *MRI system* |
| --- | --- | --- | --- | --- | --- | --- |
| 1 | Discovery^a^ | Pictures with varying disgust intensity levels | 78 (44) | 22.10 (2.68) | UESTC | 3T GE |
| 2 | Validation^a^ | Pictures with varying disgust intensity levels | 30 (16) | 21.13 (2.18) | UESTC | 3T GE |
| 3 | Modified disgust induction task dataset^a^ | Pictures with varying disgust intensity levels | 34 (15) | 23.50 (2.27) | UESTC | 3T GE |
| 4 | Chen et al.,  2021^18^ | Disgust and neutral pictures | 26 (10) | 21.73 (1.69) | Southwest University | 3T Siemens Trio |
| 5 | Chang et al., 2015^19^ | Negative and neutral pictures | 61 (Not report) | Not report | University of Colorado, Boulder | 3T Siemens Trio |
| 6 | Zhou et al., 2021^20^ | Pictures with varying fear intensity levels | 67 (34) | 21.5 (2.1) | UESTC | 3T GE |
| 7 | Gustatory dataset^a^ | Distaste liquids and neutral liquids | 30 (15) | 24.27 (2.26) | UESTC | 3T GE |
| 8 | Ultimatum game data^b^ | A series of unfairness offers | 43 (23) | 21.26  (2.12) | UESTC | 3T GE |
| 9 | Zhou et al., 2020^2^ | painful faces,  respectively non-painful controls | 238 (118) | 21.58 (2.32) | UESTC | 3T GE |
|  |  | physical pain,  respectively non-painful controls | 238 (118) | 21.58 (2.32) | UESTC | 3T GE |
| 10 | Xu et al., 2022^21^(placebo group) | Negative and neutral pictures | 30 (0) | 20.83 (2.05) | UESTC | 3T GE |
| 11 | Fastenrath et al., 2022^22, c^ | Negative and neutral pictures | 150 (75) | 22.47  (3.59) | University of Basel | 3T Siemens Verio |

^a^ current study data; ^b^ unpublished data; ^c^ we applied for a randomly selected 150 participants’ data (with gender ratio balanced) from the authors.

Supplementary Table 2. Meta-analytic decoding of the unthresholded VIDS pattern.

| *N* | *Topic* | *Corr* | *N* | *Topic* | *Corr* | *N* | *Topic* | *Corr* |
| --- | --- | --- | --- | --- | --- | --- | --- | --- |
| *1* | eating | 0.22921 | *35* | arousal | 0.16724 | *69* | negative emotional | 0.15616 |
| *2* | disgust | 0.20757 | *36* | valence | 0.16617 | *70* | symptoms | 0.15579 |
| *3* | inducing | 0.20511 | *37* | elevated | 0.16613 | *71* | illness | 0.15519 |
| *4* | food | 0.20140 | *38* | negative | 0.16609 | *72* | hyperactivity | 0.15482 |
| *5* | regulation | 0.19188 | *39* | reactivity | 0.16556 | *73* | threat | 0.15480 |
| *6* | symptom | 0.19037 | *40* | dysregulation | 0.16469 | *74* | behaviors | 0.15448 |
| *7* | anxiety | 0.18967 | *41* | monetary reward | 0.16444 | *75* | enhancing | 0.15446 |
| *8* | aversive | 0.18706 | *42* | disorders | 0.16328 | *76* | intense | 0.15435 |
| *9* | emotions | 0.18606 | *43* | pleasant | 0.16327 | *77* | anticipation | 0.15361 |
| *10* | feelings | 0.18385 | *44* | induction | 0.16290 | *78* | distress | 0.15358 |
| *11* | conditioning | 0.18282 | *45* | treatments | 0.16285 | *79* | incentive | 0.15350 |
| *12* | aberrant | 0.18251 | *46* | conditioned | 0.16266 | *80* | evaluation | 0.15307 |
| *13* | affective | 0.18175 | *47* | depressive disorder | 0.16248 | *81* | empathy | 0.15266 |
| *14* | heightened | 0.18027 | *48* | reward | 0.16220 | *82* | reactions | 0.15215 |
| *15* | negative affect | 0.18017 | *49* | loss | 0.16202 | *83* | threatening | 0.15185 |
| *16* | emotion | 0.17871 | *50* | sad | 0.16142 | *84* | regulate | 0.15167 |
| *17* | concentration | 0.17866 | *51* | emotional | 0.16131 | *85* | acute | 0.15167 |
| *18* | disorder | 0.17647 | *52* | empathic | 0.16117 | *86* | autonomic | 0.15135 |
| *19* | olfactory | 0.17549 | *53* | separation | 0.16088 | *87* | neurocognitive | 0.15135 |
| *20* | taste | 0.17472 | *54* | major depressive | 0.15971 | *88* | experiences | 0.15037 |
| *21* | neutral pictures | 0.17447 | *55* | suffering | 0.15906 | *89* | emotional valence | 0.15007 |
| *22* | emotion regulation | 0.17434 | *56* | emotional stimuli | 0.15846 | *90* | hypoactivation | 0.14969 |
| *23* | rated | 0.17373 | *57* | neutral | 0.15834 | *91* | bipolar disorder | 0.14941 |
| *24* | reward anticipation | 0.17233 | *58* | diagnosed | 0.15830 | *92* | calculated | 0.14936 |
| *25* | mood | 0.17191 | *59* | consumption | 0.15826 | *93* | depicting | 0.14935 |
| *26* | fear | 0.17152 | *60* | motivation | 0.15825 | *94* | emotional faces | 0.14901 |
| *27* | pictures | 0.17145 | *61* | experienced | 0.15821 | *95* | disruption | 0.14884 |
| *28* | salience | 0.17088 | *62* | positive | 0.15795 | *96* | affect | 0.14800 |
| *29* | experiencing | 0.17023 | *63* | health | 0.15790 | *97* | severity | 0.14755 |
| *30* | stress | 0.16999 | *64* | emotional information | 0.15767 | *98* | pain | 0.14753 |
| *31* | salience network | 0.16910 | *65* | dysfunctional | 0.15729 | *99* | reported | 0.14677 |
| *32* | negative emotions | 0.16892 | *66* | motivational | 0.15677 | *100* | salient | 0.14653 |
| *33* | ratings | 0.16752 | *67* | unpleasant | 0.15659 |  |  |  |
| *34* | emotionally | 0.16729 | *68* | subjective | 0.15646 |  |  |  |

Supplementary Table 3. Prediction-outcome correlations (mean±std) with various numbers of voxels (for the Discovery Cohort).

| Number of voxels | | Vis | | SM | | dA | | vA | | Limb | | FP | | DMN | | Cons | | subC | Whole-brain |
| --- | --- | --- | --- | --- | --- | --- | --- | --- | --- | --- | --- | --- | --- | --- | --- | --- | --- | --- | --- |
| 50 | 0.153  (0.038) | | 0.308  (0.050) | | 0.230  (0.048) | | 0.379  (0.037) | | 0.283  (0.045) | | 0.293  (0.045) | | 0.346  (0.046) | | 0.347  (0.051) | | 0.359  (0.042) | | 0.326  (0.055) |
| 150 | 0.213  (0.039) | | 0.322  (0.041) | | 0.272  (0.038) | | 0.390  (0.036) | | 0.329  (0.040) | | 0.325  (0.042) | | 0.380  (0.044) | | 0.365  (0.044) | | 0.399  (0.034) | | 0.361  (0.046) |
| 250 | 0.238  (0.031) | | 0.305  (0.034) | | 0.277  (0.031) | | 0.391  (0.034) | | 0.350  (0.035) | | 0.328  (0.037) | | 0.382  (0.040) | | 0.354  (0.036) | | 0.397  (0.033) | | 0.357  (0.041) |
| 500 | 0.239  (0.025) | | 0.281  (0.027) | | 0.255  (0.023) | | 0.370  (0.026) | | 0.355  (0.028) | | 0.327  (0.030) | | 0.373  (0.032) | | 0.337  (0.032) | | 0.382  (0.026) | | 0.353  (0.038) |
| 750 | 0.239*  (0.021) | | 0.275  (0.024) | | 0.243*  (0.020) | | 0.346  (0.021) | | 0.346  (0.024) | | 0.319  (0.025) | | 0.376  (0.029) | | 0.366  (0.031) | | 0.368  (0.022) | | 0.418  (0.034) |
| 1000 | 0.240*  (0.018) | | 0.271*  (0.021) | | 0.234*  (0.018) | | 0.334  (0.019) | | 0.336  (0.022) | | 0.318  (0.023) | | 0.389  (0.026) | | 0.389  (0.026) | | 0.358  (0.020) | | 0.454  (0.033) |
| 2000 | 0.258*  (0.013) | | 0.275*  (0.014) | | 0.244*  (0.014) | | 0.325*  (0.014) | | 0.330*  (0.017) | | 0.347*  (0.017) | | 0.422  (0.018) | | 0.416  (0.019) | | 0.347*  (0.014) | | 0.512  (0.026) |
| 3500 | 0.267*  (0.010) | | 0.280*  (0.010) | | 0.251*  (0.010) | | 0.334*  (0.009) | | 0.345*  (0.012) | | 0.359*  (0.011) | | 0.436  (0.013) | | 0.428  (0.013) | | 0.336*  (0.010) | | 0.534  (0.018) |
| 6500 | 0.272*  (0.006) | | 0.283*  (0.007) | | 0.255*  (0.006) | | 0.339*  (0.005) | | 0.352*  (0.007) | | 0.366*  (0.007) | | 0.443*  (0.008) | | 0.433*  (0.009) | | 0.332*  (0.005) | | 0.548  (0.013) |
| 10000 | 0.274*  (0.005) | | 0.284*  (0.004) | | 0.257*  (0.004) | | 0.341*  (0.002) | | 0.355*  (0.004) | | 0.368*  (0.005) | | 0.446*  (0.006) | | 0.435*  (0.007) | | 0.332*  (0.001) | | 0.553  (0.011) |
| 14000 | NA | | NA | | NA | | NA | | NA | | NA | | NA | | 0.436*  (0.006) | | NA | | 0.557  (0.009) |
| 30000 | NA | | NA | | NA | | NA | | NA | | NA | | NA | | 0.438*  (0.003) | | NA | | 0.560  (0.006) |
| 65000 | NA | | NA | | NA | | NA | | NA | | NA | | NA | | NA | | NA | | 0.562  (0.004) |
| 150000 | NA | | NA | | NA | | NA | | NA | | NA | | NA | | NA | | NA | | 0.563  (0.002) |
| Full | 0.276 | | 0.285 | | 0.258 | | 0.342 | | 0.356 | | 0.370 | | 0.449 | | 0.438 | | 0.332 | | 0.563 |

Vis, visual network; SM, somatomotor network; dA, dorsal attention network; vA, ventral attention network; Limb, limbic network; FP, frontoparietal network; DMN, default model network; Cons, consciousness network; subC, subcortical network; NA, not applicable. *indicates that the prediction is significant lower as compared with the whole-brain model using the same number of voxels (two-tailed *Z*-test; *P*<0.05, Bonferroni corrected).

Supplementary Table 4. Comparing prediction (correlation) of VIDS, PINES and VIFS.

| **Datasets** | **VIDS** | **PINES** | **VIFS** |
| --- | --- | --- | --- |
| Discovery | 0.56 [0.49, 0.62]; 0.88±0.01^a^ | 0.24 [0.13, 0.33]; 0.45±0.05 | 0.35 [0.24, 0.43]; 0.75±0.04 |
| Validation | 0.62 [0.53, 0.70]; 0.89±0.03 | 0.30 [0.14, 0.44]; 0.50±0.08 | 0.32 [0.17, 0.45]; 0.68±0.07 |
| VIFS discovery | 0.39 [0.29, 0.47]; 0.64±0.05 | 0.38 [0.28, 0.47]; 0.59±0.04 | 0.57 [0.49, 0.63]; 0.89±0.01^a^ |
| PINES holdout | 0.23 [0.11, 0.33]; 0.45±0.07 | 0.72 [0.65, 0.77]; 0.90±0.01 | 0.29 [0.17, 0.38]; 0.63±0.04 |

We applied the VIDS, PINES, and VIFS to subjective disgust, fear discovery, and general negative emotion holdout datasets and calculated the overall (bootstrapped 95% CI) as well as within-subject (mean±SE) prediction−outcome correlations between the pattern expressions and the true ratings.

^a^ indicates cross-validated.

Supplementary Table 5. The 20 distributions of 100 RMB used in Study 8.

| Proposer | Responder (The participant) | Unfairness level |
| --- | --- | --- |
| 97.5 | 2.5 | **Unfairness level 5** |
| 95 | 5 |  |
| 92.5 | 7.5 |  |
| 90 | 10 |  |
| 87.5 | 12.5 | **Unfairness level 4** |
| 85 | 15 |  |
| 82.5 | 17.5 |  |
| 80 | 20 |  |
| 77.5 | 22.5 | **Unfairness level 3** |
| 75 | 25 |  |
| 72.5 | 27.5 |  |
| 70 | 30 |  |
| 67.5 | 32.5 | **Unfairness level 2** |
| 65 | 35 |  |
| 62.5 | 37.5 |  |
| 60 | 40 |  |
| 57.5 | 42.5 | **Unfairness level 1** |
| 55 | 45 |  |
| 52.5 | 47.5 |  |
| 50 | 50 |  |
